# ARID1A regulates histone octamer transfer activity of human canonical BAF complex

**DOI:** 10.1101/2025.05.21.655315

**Authors:** Naoe Moro, Yukiko Fujisawa-Tanaka, Shinya Watanabe

## Abstract

Mutations that impact subunits of mammalian SWI/SNF (mSWI/SNF or BAF) chromatin remodeling complexes are found in over 20% of human cancers. Among these subunits, ARID1A is the most frequently mutated gene, occurring in over 8% of various cancers. The majority of ARID1A mutations are frameshift or nonsense mutations, causing loss of function. Previous studies have suggested that ARID1A may facilitate interactions between BAF complexes and various transcriptional coactivators, but a biochemical role for ARID1A in BAF remodeling activity has not been identified. Here, we describe the in vitro reconstitution of the cBAF, PBAF, and ncBAF complexes, and we compare their biochemical activities. In addition, we reconstitute a variety of cBAF subcomplexes, defining roles for several subunits in high affinity nucleosome binding and nucleosome sliding activity. Remarkably, we find that the ARID1A subunit of cBAF is largely dispensable for nucleosome binding, nucleosome sliding, and ATPase activity, but ARID1A is required for cBAF to transfer histone octamers between DNA templates. These data suggest a model in which the histone octamer transfer activity of BAF complexes is key for cancer prevention.

## INTRODUCTION

ATP-dependent chromatin remodeling enzymes are involved in almost all DNA metabolic processes, such as transcription, DNA repair, and DNA replication. They utilize the energy from ATP hydrolysis to directly alter chromatin structure by catalyzing the sliding of histone octamers *in cis* along DNA, the transfer or exchange of histone H2A/H2B dimers, or the ejection or transfer of an entire histone octamer (1). How each of these distinct biochemical activities contributes to a specific cellular function is not clear since inactivation of a key remodeling subunit typically inactivates all activities of a remodeling enzyme.

Genes encoding subunits of mammalian SWI/SNF (mSWI/SNF or BAF) chromatin remodeling complexes are mutated in over 20% of human cancers (2,3). Mutations impacting BAF subunits also cause neurodevelopment disorders, such as Coffin-Siris syndrome (4). Human BAF complexes exist as three major complexes -- cBAF (canonical BRG1/BRM-associated factors), PBAF (Polybromo-associated BAF), and ncBAF (non-canonical BAF), which differ based on their distinct subunit compositions. To date, the cBAF complex is believed to be composed of 12 subunits, 13 subunits for PBAF, and 10 subunits for the ncBAF complex. Several BAF subunits also have multiple isoforms, extending the diversity of BAF complex composition. CryoEM analyses of cBAF and PBAF indicate that these complexes consist of three modules – an ATPase module, an ARP module, and a Base module (5,6). The ATPase module is composed of the ATPase domain of the BRG1 (SMARCA4) subunit. The ARP module contains ß-actin and BAF53a (ACTL6A) that are associated with the N-terminal HSA domain of BRG1, and the ARP module hinges the ATPase and Base modules. The Base module consists of 5 core subunits shared among both cBAF and PBAF complexes (BAF170 (SMARCC1), BAF155 (SMARCC2), BAF60 (SMARCD1), BAF57 (SMARCE1), and BAF47 (SMARCB1)), as well as cBAF- or PBAF-specific subunits. cBAF and PBAF engage the nucleosome by sandwiching it between the ATPase module and the BAF47 (SMARCB1) subunit within the Base module.

The AT-Rich Interactive Domain-containing protein 1A (ARID1A) is a cBAF-specific subunit and a component of the Base module. ARID1A is the most frequently mutated gene among BAF subunits (∼8% of all types of cancer) (7). In gynecologic cancers, ARID1A mutations are found in 60% of ovarian clear-cell carcinomas (OCCC), 30% of ovarian endometrioid carcinoma (OEC), and 40% of low-grade endometrioid adenocarcinomas (8,9). The majority of ARID1A mutations are frameshift or nonsense mutations. Because the C-terminal domain of ARID1A is required for assembly into cBAF, all truncating mutations cause loss of function. Notably, ARID1A has a mutually exclusive paralog subunit, ARID1B, which was identified as a major vulnerability in ARID1A-deficient cancer cell lines (10), suggesting that ARID1B is a therapeutic target in ARID1A-deficient cancers. How ARID1A or ARID1B contributes to the biochemical functions of cBAF is still largely unknown.

Here, we describe the reconstitution of all three major BAF complexes, cBAF, PBAF, and ncBAF, and we perform the direct comparison of their biochemical activities. In addition, we reconstitute a variety of cBAF subcomplexes to define roles for several subunits in complex integrity, high affinity nucleosome binding, and nucleosome sliding activity. Remarkably, we find that ARID1A is required for the histone octamer transfer activity of cBAF, but loss of ARID1A has little effect on other biochemical activities, including ATPase, nucleosome binding, and nucleosome sliding. We also find that ARID1B, a mutually exclusive paralog of ARID1A, can substitute for ARID1A, demonstrating that ARID1B is also required for histone octamer transfer activity. Our study reveals the biochemical function of ARID1A/ARID1B in BAF-mediated chromatin remodeling and suggests a model in which dysregulation of histone octamer transfer activity of BAF complexes may lead for cancer formation.

## RESULTS

### Histone octamer transfer activity is conserved in all three BAF complexes

To investigate the detailed biochemical properties of the three major BAF complexes, we reconstituted human cBAF, PBAF, and ncBAF complexes with the full-length, recombinant subunits using a baculoviral expression system and Strep-tag affinity purification. The subunit compositions were confirmed by SDS-PAGE (**Figure 1A**) and mass-spectrometry (Supplemental table S1). The individual BRG1 ATPase subunit was also purified, so that its activity could be compared to the intact BAF complexes (**Figure 2A**).

**Figure 1.**
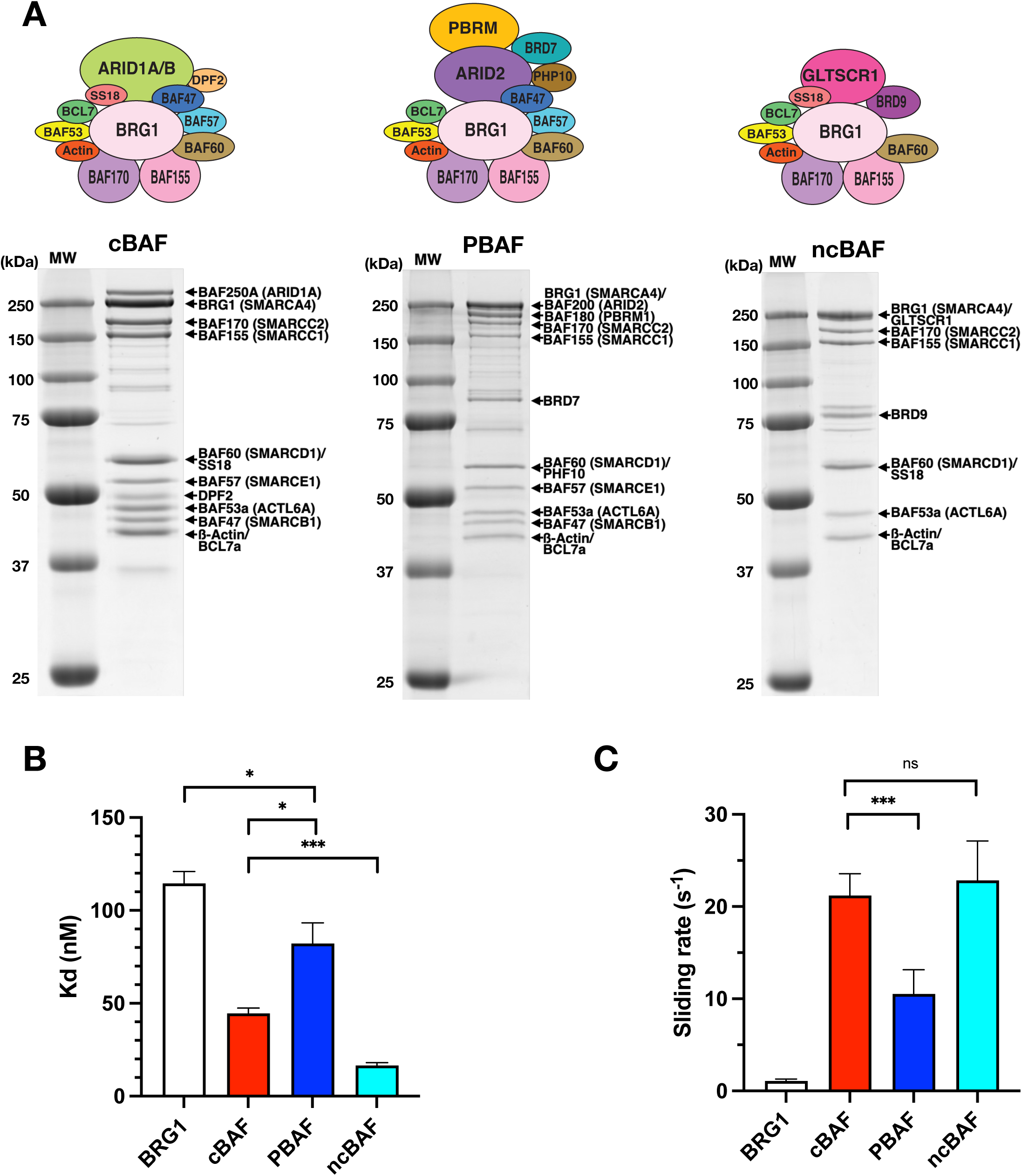
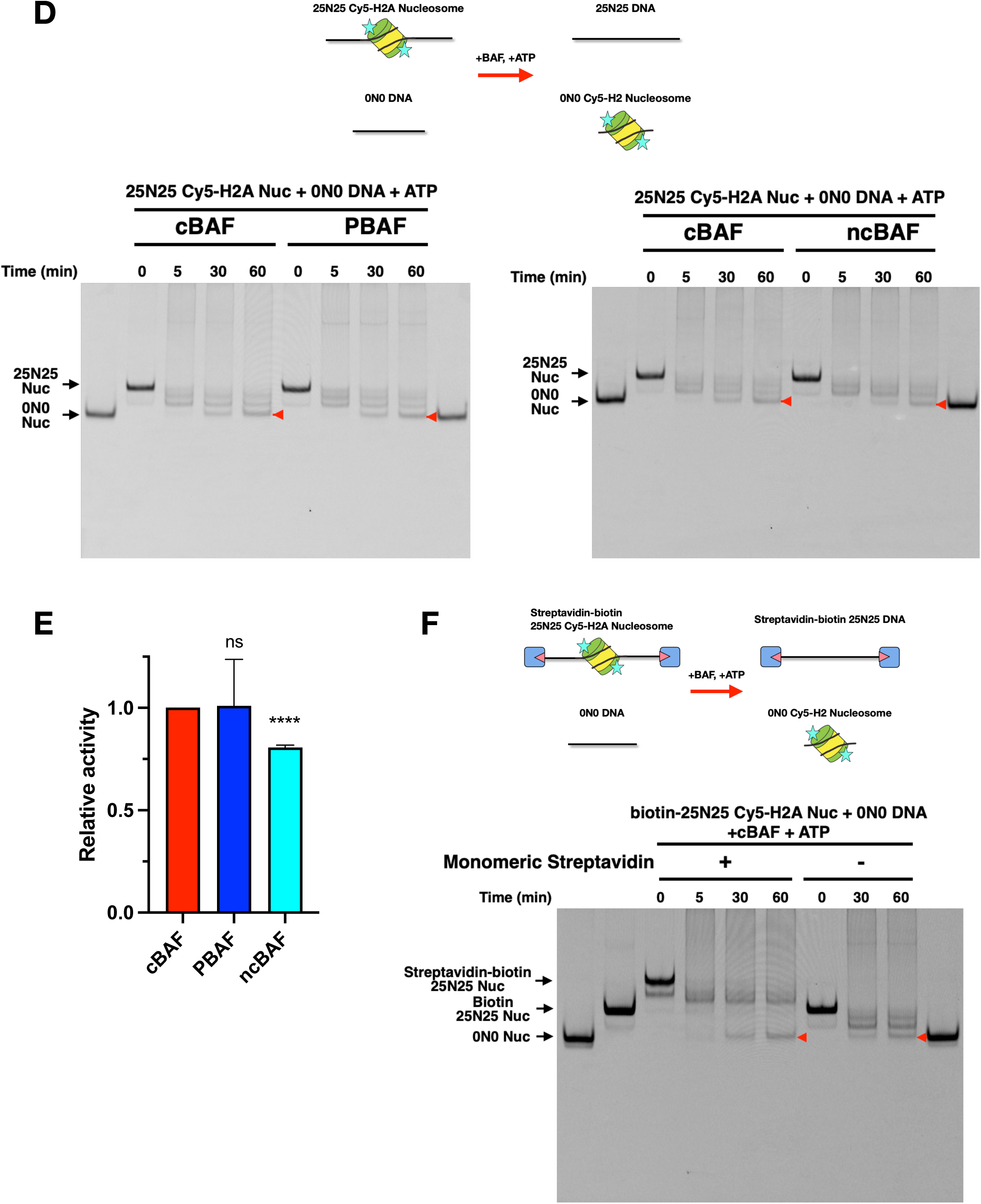
Histone octamer transfer activity is conserved in all three BAF complexes. (A) Coomassie-staining SDS-PAGE gels of purified cBAF (Left panel), PBAF (Middle panel), and ncBAF (Right panel). (B) Nucleosome-binding affinity of cBAF, PBAF, and ncBAF. (C) Nucleosome sliding activity of cBAF, PBAF, and ncBAF. (D) Histone octamer transfer activity of cBAF, PBAF, and ncBAF. Schematic of histone octamer transfer assay (Top panel). Representative Cy5-scanned native gels of histone octamer transfer assay (Bottom panels). Red arrow indicates 0N0 nucleosomes produced by octamer transfer. (E) Quantification of histone octamer transfer activity. Each activity was normalized to cBAF activity. (F) Representative Cy5-scanned native gel of histone octamer transfer assay using streptavidin-biotinylated nucleosomes. For Fig. 1B, 1C, and 1E, each error bar represents the standard error from at least three independent experiments using at least two independent BAF preparations. ****P<0.0001, ***P<0.001, *P<0.05; ns, not significant.

**Figure 2.**
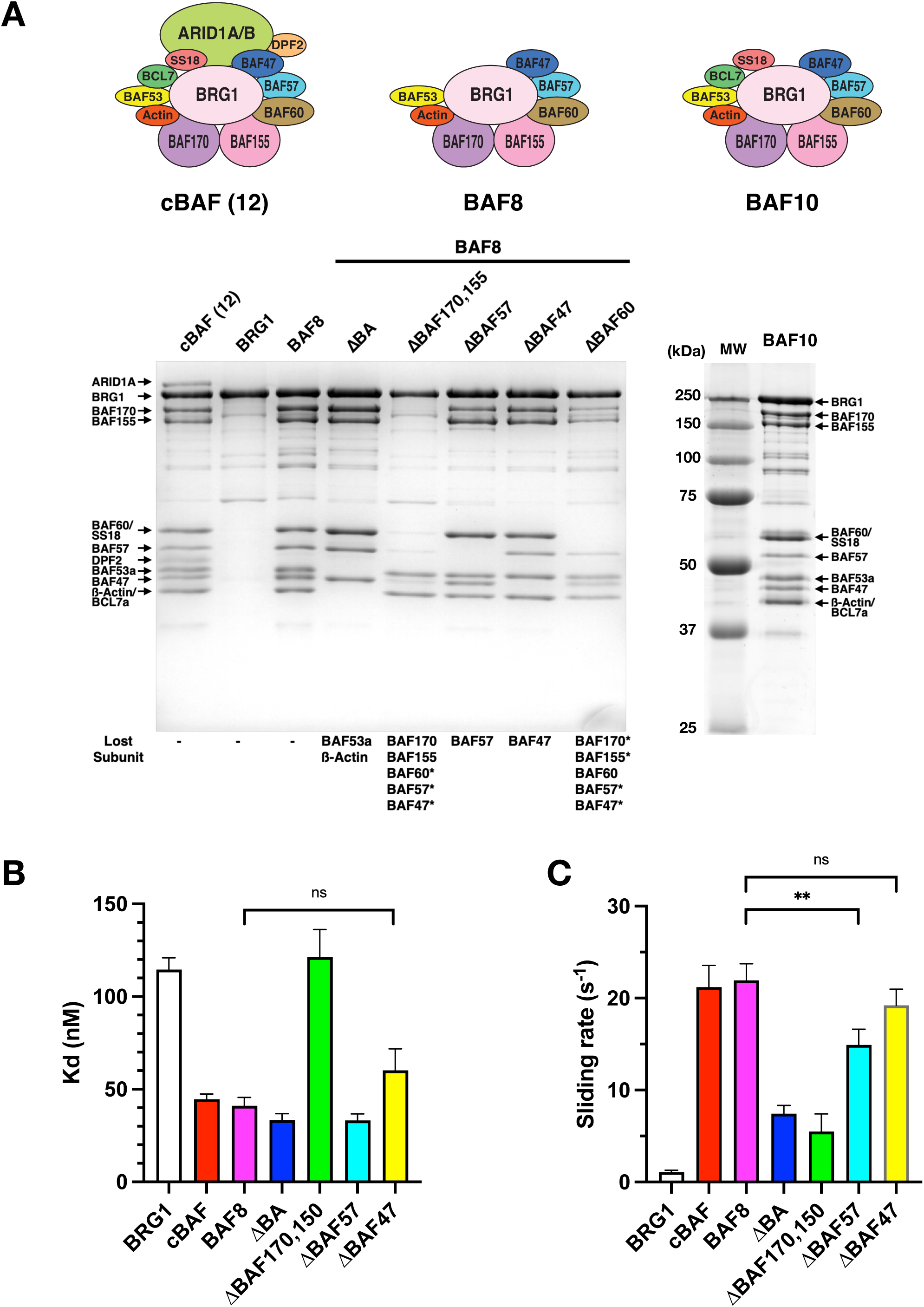
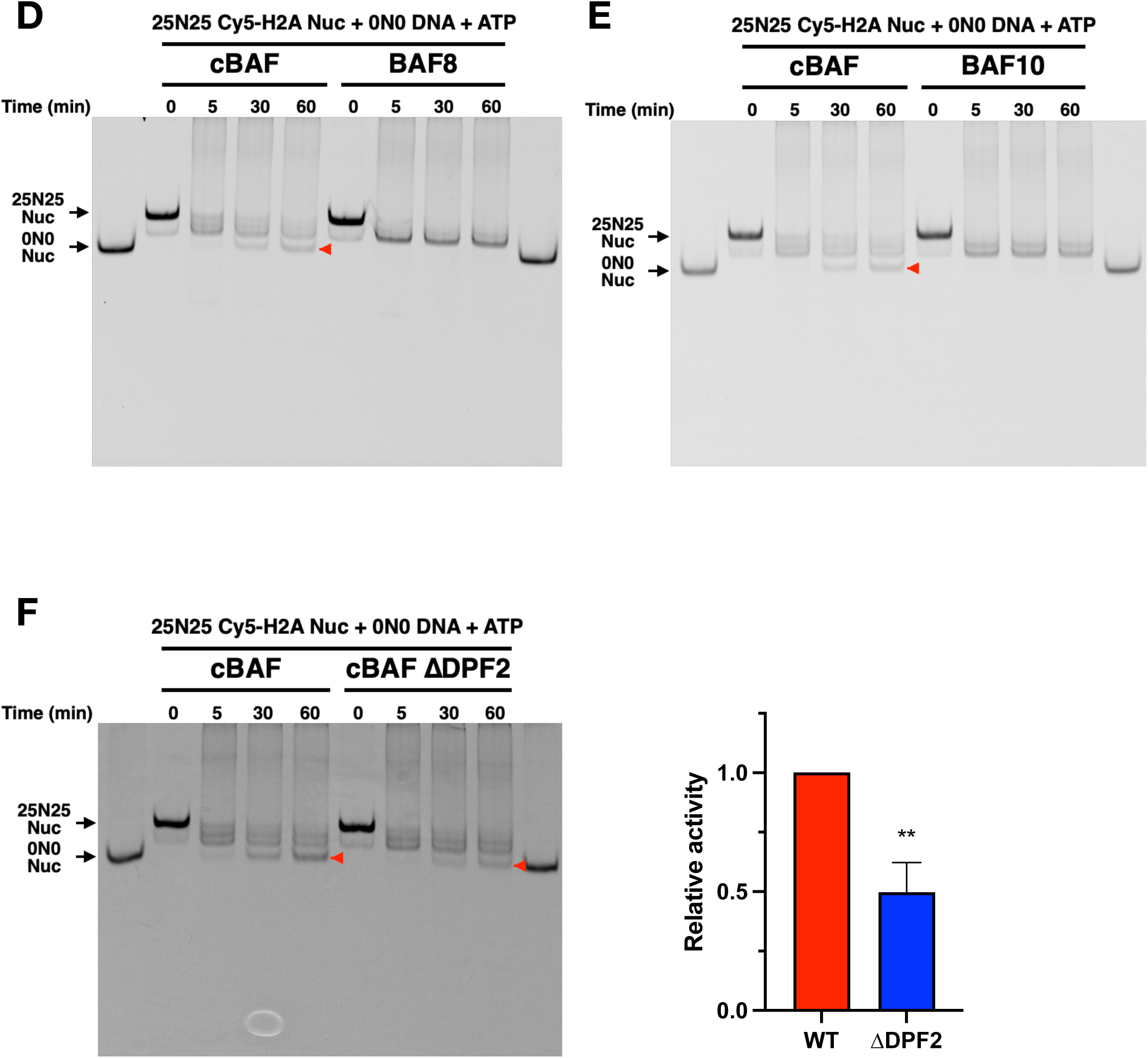
The core 8-subunit BAF complex lacks histone octamer transfer activity. (A) Coomassie-staining SDS-PAGE gels of purified BAF subcomplexes. (B) Nucleosome-binding affinity of BAF8 subcomplexes. (C) Nucleosome sliding activity of BAF8 subcomplexes. (D) Representative Cy5-scanned native gel of histone octamer transfer assay for BAF8. Red arrow indicates 0N0 nucleosomes produced by octamer transfer. (E) Representative Cy5-scanned native gel of histone octamer transfer assay for BAF10. (F) Representative Cy5-scanned native gel (Left panel) and quantification (Right panel) of histone octamer transfer assay for cBAF ΔDPF2 complex. The activity was normalized to cBAF WT activity. For Fig. 2B, 2C, 2F, each error bar represents the standard error from at least three independent experiments using at least two independent BAF preparations. **P<0.01; ns, not significant.

To quantify the ability of each BAF complex to bind its nucleosomal substrate, a fluorescence polarization (FP) assay was employed (**Figures 1B and S1A**). Recombinant nucleosomes were reconstituted with human histone octamers that contained a Cy5-labelled H2A subunit and a 197 bp DNA template that harbored a ‘601’ nucleosome positioning sequence. As expected, the BRG1 subunit bound to nucleosomes weaker, in comparison to the BAF complexes, with a Kd of 115 nM. The ncBAF complex exhibited the highest affinity for nucleosomes, with a Kd of 17 nM, followed by cBAF at 45 nM and PBAF at 82nM.

The differences in binding affinity were paralleled by the ATP-dependent nucleosome sliding activity of each remodeler. Using a quantitative restriction enzyme accessibility assay, each BAF complex catalyzed the sliding of nucleosomes at rates that were at least 10-fold higher than the isolated BRG1 subunit (**Figures 1C, S1B, and S2B**). Furthermore, and consistent with their differing binding affinities, the cBAF and ncBAF complexes were more effective than PBAF in this sliding assay (**Figures 1C and S1B**). The decreased sliding rate of PBAF, compared to the activities of cBAF and ncBAF, is consistent with previously published results that used endogenous BAF complexes and a modified nucleosome library (11).

Whereas most remodeling enzymes can slide nucleosomes in cis, SWI/SNF complexes are also able to transfer an entire histone octamer from one nucleosome to a different DNA fragment. To measure this histone octamer transfer activity, BAF complexes were incubated with a Cy5-labelled histone H2A 25N25 nucleosome and a 147 bp DNA (0N0), and the reaction was initiated by addition of ATP (**Figure 1D, Top**). The histone octamer transfer activity of BAF led to the ATP-dependent formation of a Cy5-labelled 0N0 nucleosome, monitored by Native-PAGE and a Cy5 scan (see red arrowhead). Note that formation of this product requires addition of the 0N0 DNA (**Figure S1C**). All three BAF complexes showed comparable histone octamer transfer activity (**Figures 1D, Bottom, and 1E**). Notably, under these assay conditions, almost all nucleosomes were re-positioned from the original center position to the end of the 25N25 DNA fragment within 5 minutes (note formation of faster migrating species). The product of the histone octamer transfer reaction, the 0N0 nucleosome, accumulated at 30-60 minutes, suggesting that the histone octamer transfer reaction may not be coupled to nucleosome sliding.

One possibility is that histone octamer transfer activity involves the sliding of histone octamers “off the ends” of the original DNA fragment. To test this idea, both ends of the 25N25 DNA fragment were modified with biotin, so that streptavidin could be used to block each DNA end, as previously demonstrated (12) **(Figure 1F, Top)**. Cy5-H2A-containing nucleosomes were reconstituted with the biotinylated 25N25 DNA fragments, and the addition of monomeric streptavidin led to the expected mobility shift of the biotin-25N25 nucleosomes on Native-PAGE (**compare lane 2 and 3 in Figure 1F, Bottom**). We then compared the histone octamer transfer activity of cBAF in the presence or absence of monomeric streptavidin (**compare lane 6 and 9 in Figure 1F**). Importantly, the addition of streptavidin did not affect the histone transfer activity of cBAF, suggesting that a free DNA end is not required for histone octamer transfer.

It is also possible that BAF complexes do not transfer histone octamers, but rather BAF can only evict histone octamers and that these dissociated histones are then passively assembled onto free DNA. To assess the histone eviction activity of BAF complexes, nucleosomes were reconstituted onto a Cy5-lebelled 25N25 DNA fragment and incubated with BAF complexes in the absence of 0N0 DNA. In this assay, histone octamer eviction leads to an increased amount of free Cy5-lebelled 25N25 DNA, monitored by Native-PAGE and a Cy5 scan. All three BAF complexes increased the fraction of free 25N25 DNA by only ∼5% within 5 min after ATP addition, and this level remained unchanged for the extended time course to 90 min (**Figure S1D**). These results suggest that a limited amount of histone octamer eviction may be coupled to nucleosome sliding, unlike histone octamer transfer. Importantly, this result suggests that histone octamer eviction activity is not a major contributor to histone octamer transfer activity.

### The core 8-subunit BAF complex lacks histone octamer transfer activity

To further investigate the assembly and function of BAF complexes, we reconstituted a BAF subcomplex with 8 core subunits shared between cBAF and PBAF complexes (BRG1, BAF170, BAF155, BAF60, BAF57, BAF47, BAF53a, and ß-Actin), which we refer to here as “BAF8” (**Figure 2A, BAF8**). Various subunits were also omitted from the BAF8 reconstitution, leading to a set of smaller subcomplexes (**Figure 2A**). Previous glycerol gradient purification analyses of endogenous BAF complexes suggested that there may be a BAF core module containing 5 subunits (BAF170, BAF155, BAF60, BAF57, and BAF47) that stably exists as a precursor (13). When BAF53a and ß-Actin were removed from BAF8, the assembly of other subunits with BRG1 was not impacted (**Figure 2A, ΔBA**). Since endogenous ß-Actin exists abundantly, this result suggests that BAF53a might be required for the association of ß-Actin with the BAF complex. Since BAF170 and BAF155 are interchangeable (5), both were eliminated from BAF8, leading to the loss of many subunits, with the exception of ß-Actin and BAF53a (**Figure 2A, ΔBAF170,155**). Removal of either BAF57 or BAF47 had no effect on the association of the remaining subunits with the complex (**Figure 2A, ΔBAF57 and ΔBAF47**), whereas elimination of BAF60 resulted in reduced amounts of all Base subunits (BAF170, BAF155, BAF57, and BAF47) (**Figure 2A, ΔBAF60**). These results suggest that (1) the ARP module and the Base module interact independently with BRG1; (2) BAF170, BAF155, and BAF60 are required for the overall integrity of the Base module, consistent with previous work with endogenous complexes (13).

We next measured the nucleosome-binding affinity and nucleosome sliding activity of BAF8 and BAF8 subcomplexes (**Figures 2B, 2C, S2A, and S2B)**. The cBAF and BAF8 complexes were nearly identical in these assays, suggesting that the 4 other subunits of cBAF (BCL7, SS18, ARID1A, and DPF2) do not contribute significantly to nucleosome binding or nucleosome sliding activities. Likewise, the ΔBAF57 and ΔBAF47 subcomplexes had similar nucleosome sliding rates and nucleosome binding affinities as compared to the BAF8 and cBAF complexes, indicating that BAF57 and BAF47 also do not contribute significantly to these activities, despite the fact that BAF47 is known to make contact with the nucleosomal acidic patch within the cBAF-nucleosome and PBAF-nucleosome complexes (5,6). In contrast, loss of the ARP module (ΔBA) reduced the nucleosome sliding rate, though nucleosome binding affinity was unaffected. Strikingly, loss of the Base module (ΔBAF170,155) reduced both nucleosome sliding activity and nucleosome binding, with binding affinity similar to that of the isolated BRG1 subunit. These results suggest that (1) the ARP module is required for proper nucleosome sliding, but not for nucleosome-binding; (2) the Base module subunits, except for BAF60 and BAF47, are required for proper nucleosome sliding and nucleosome-binding.

### ARID1A is required for histone octamer transfer activity of cBAF

Finally, we measured the histone octamer transfer activity of BAF8. Surprisingly, BAF8 did not exhibit detectable histone octamer transfer activity (**Figure 2D**), suggesting that the four subunits of cBAF that are missing from BAF8 (BCL7a, SS18, ARID1A, and DPF2) are required for histone octamer transfer activity. To identify which subunit(s) is required for histone octamer transfer activity, we added the BCL7a and SS18 subunits to BAF8, leading to a BAF10 subcomplex (**Figure 2A**). However, BAF10 remained inactive in the histone octamer transfer assay (**Figure 2E**). Reconstitution of a cBAF complex that only lacked the DPF2 subunit had no effect on the overall integrity of cBAF (**Figure S2C**), but reduced histone octamer transfer activity (**Fig. 2F**), suggesting that DPF2 contributes to histone octamer transfer activity, but is not essential. Finally, we assembled a cBAF complex that lacks ARID1A (ΔARID1 complex). Removal of ARID1A did not grossly change the composition of BAF (**Figure 3A**), except for reduced level of DPF2, consistent with the known interaction between ARID1A and DPF2 (5). Whereas loss of ARID1A led only to small defects in ATPase, nucleosome binding, and nucleosome sliding activities (**Figures 3B-3D**, **S3A, and S3B)**, the ΔARID1 complex was inactive in the histone octamer transfer assay (**Figure 3E**). Adding back purified, recombinant ARID1A (**Figure S3C**) to the ΔARID1 complex restored histone octamer transfer activity (**Figure 3F**), confirming that ARID1A is required for the histone octamer transfer activity of cBAF.

**Figure 3.**
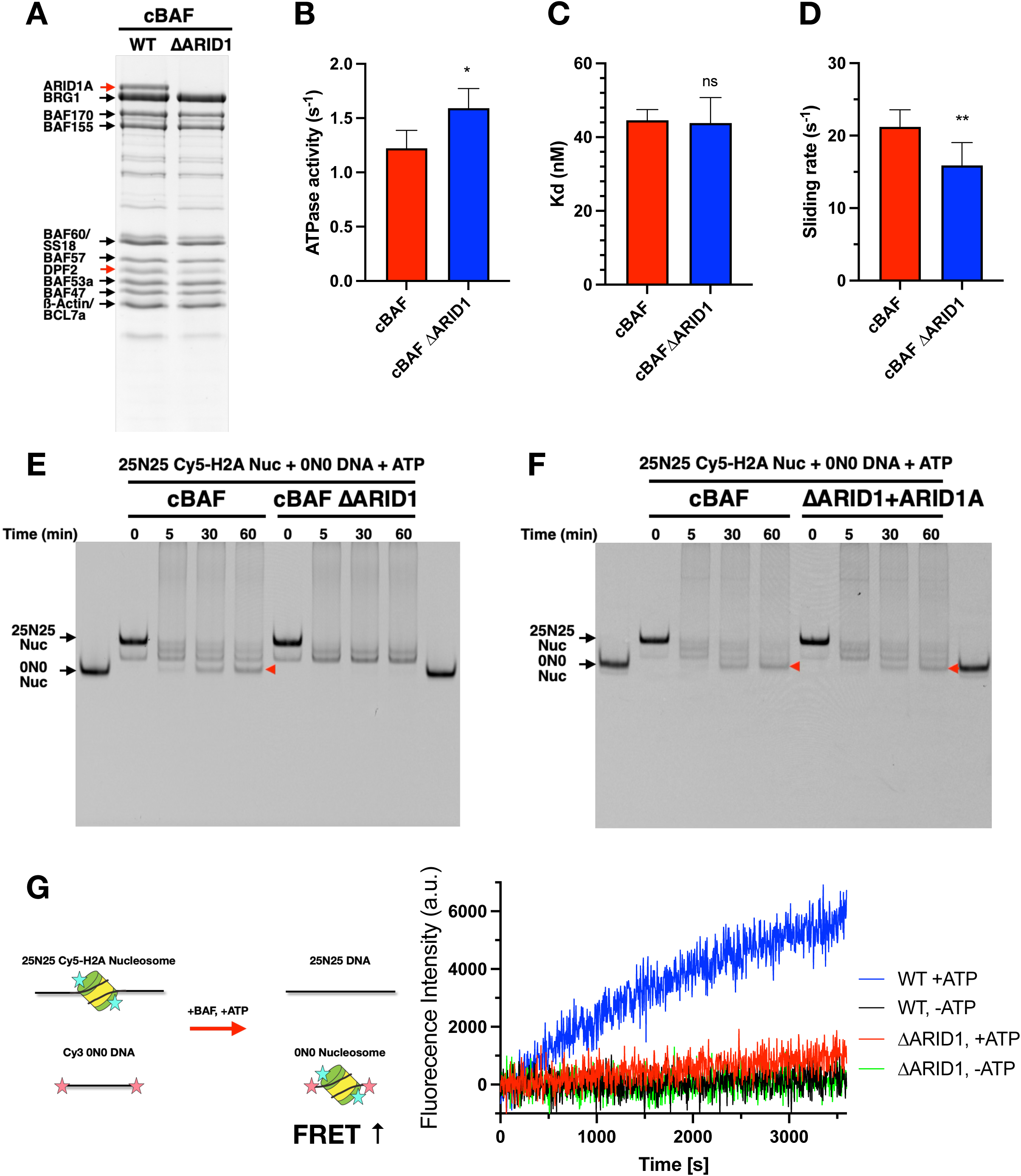
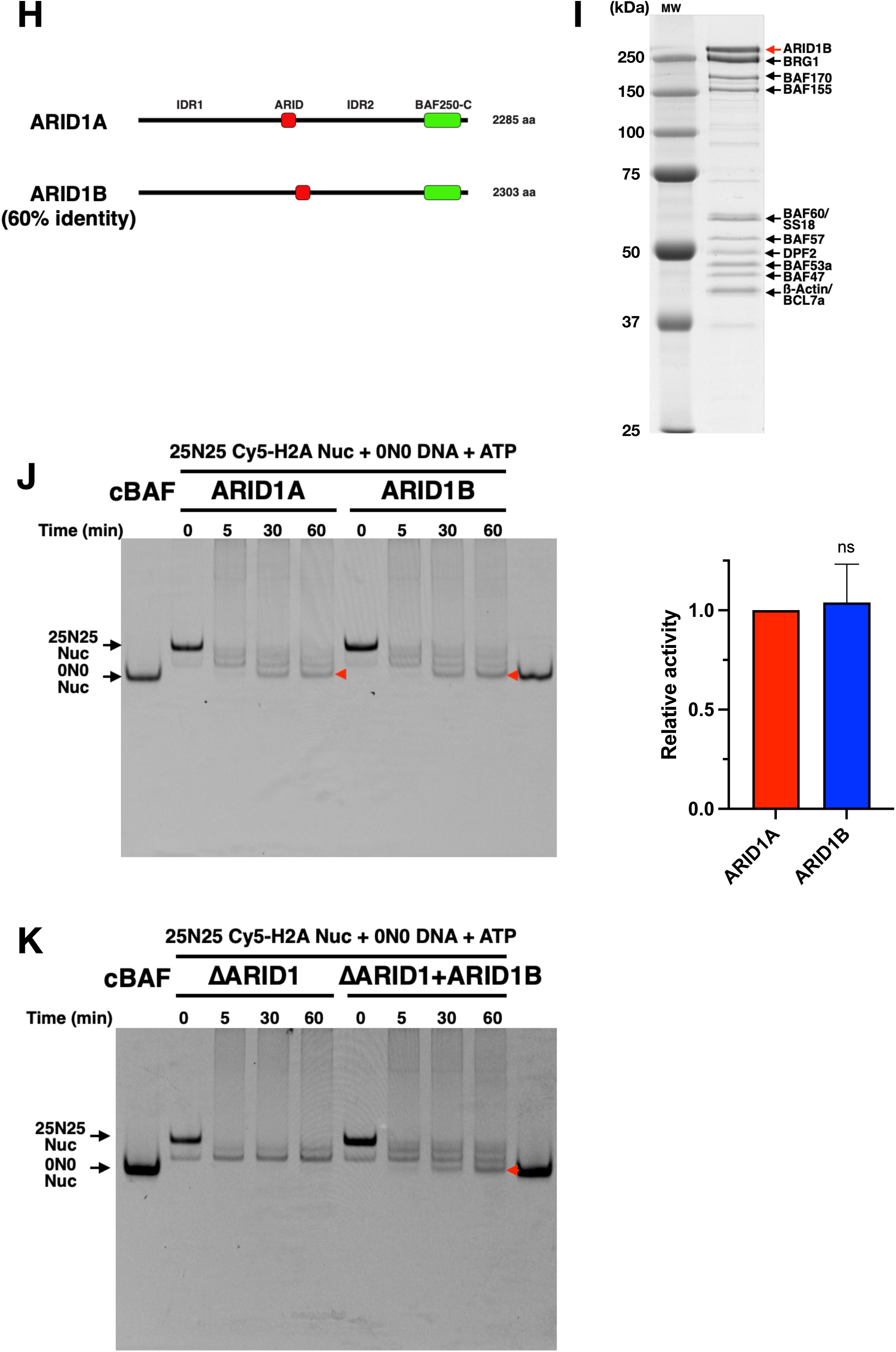
ARID1A/ARID1B are required for histone octamer transfer activity. (A) Coomassie-staining SDS-PAGE gel of purified cBAF ΔARID1 complex. (B) ATPase activity of cBAF ΔARID1. (C) Nucleosome-binding affinity of cBAF ΔARID1. (D) Nucleosome sliding activity of cBAF ΔARID1. (E) Representative Cy5-scanned native gel of histone octamer transfer assay for cBAF ΔARID1. Red arrow indicates 0N0 nucleosomes produced by octamer transfer. (F) Representative Cy5-scanned native gel of histone octamer transfer assay for cBAF ΔARID1 after adding back ARID1A. (G) FRET-based assay for monitoring histone octamer transfer activity. Schematic of the FRET-based assay (Left panel). Representative FRET signal changes of cBAF WT and cBAF ΔARID1 in the presence or absence of ATP (Right panel). (H) Domain structures of ARID1A and ARID1B. (I) Coomassie-staining SDS-PAGE gel of purified ARID1B-containing cBAF. (J) Representative Cy5-scanned native gel of histone octamer transfer assay for ARID1B-containing cBAF. The activity was normalized to ARID1A-containing cBAF activity. (K) Representative Cy5-scanned native gel of histone octamer transfer assay for cBAF ΔARID1 after adding back ARID1B. For Fig. 3B, 3C, 3D, and 3J, each error bar represents the standard error from at least three independent experiments using at least two independent BAF preparations. **P<0.01, *P<0.05; ns, not significant.

In order to monitor histone octamer transfer activity by an alternative method, we developed a FRET-based assay. In this assay, BAF complexes were incubated with Cy5-labelled histone H2A nucleosomes and a Cy3-labelled DNA fragment. Transfer of the Cy5-labelled octamer (donor fluorophore) to the Cy3-labelled DNA (acceptor fluorophore) is expected to lead to an increase in FRET (**Figure 3G, Left**). As expected, in the presence of cBAF, an ATP-dependent increase in FRET signal was observed over time (**Figure 3G, Right, Blue line**). On the other hand, the ΔARID1 complex was nearly inactive in this FRET-based histone octamer transfer assay (**Figure 3G, Right, Red line**). Taking together, these findings demonstrate that ARID1A is required for the histone octamer transfer activity of cBAF.

### ARID1B is also required for the histone octamer transfer activity of cBAF

ARID1B is a mutually exclusive paralog subunit of ARID1A with 60% identity (**Figure 3H**). Loss of ARID1B is synthetically lethal in ARID1A-deficient cancer cells, suggesting that ARID1B is a therapeutic target in ARID1A-deficient cancers. To test whether ARID1B can promote the histone octamer transfer activity of cBAF, we reconstituted an ARID1B-containing cBAF complex. The subunit composition of this cBAF complex was the same as that of ARID1A-containing cBAF (**Figure 3I**). Furthermore, the histone octamer transfer assay revealed that the ARID1B-containing complex exhibited comparable histone octamer transfer activity to the ARID1A-containing complex (**Figure 3J**), demonstrating that ARID1B can also support the histone octamer transfer activity of cBAF. Furthermore, we purified recombinant ARID1B (**Figure S3D**) and added it back to the reaction in the presence of cBAF ΔARID1 complex. Similar to ARID1A, add-back of ARID1B to the ΔARID1 complex restored histone octamer transfer activity. Taking together, these results revealed that ARID1A and ARID1B are key subunits required for the histone octamer transfer activity of cBAF.

### A part of IDR2 is required for histone octamer transfer activity of cBAF

Previous studies have identified four distinct domains within ARID1A (**Figure 4A**). The C-terminus of ARID1A is highly conserved between the two ARID1 paralogs, and this region contains 7 ARM domains that are key for assembly of ARID1 subunits into the cBAF complex. There are two intrinsically disordered regions (IDR1 and IDR2) that can form phase condensates in vitro and contribute to ARID1A function in vivo (14). Finally, there is an ARID domain that confers DNA binding to AT-rich DNA sequences (**Figure 4A**). To identify a region in ARID1A required for the histone octamer transfer activity of cBAF, we initially expressed and reconstituted cBAF complexes with three different N-terminal truncation derivatives of ARID1A (**Figure 4A**). A1A Δ990 removes IDR1, A1A Δ1124 removes both IDR1 and the ARID domain, and A1A Δ1960 removes IDR1, ARID, and IDR2 (**Figure 4A**). Each of these ARID1A truncation derivatives assembled into cBAF complexes, as confirmed by SDS-PAGE and western blot (**Figure 4B**). It should be noted that the levels of DPF2 were reduced in the cBAF complex that contains the largest truncation, A1A Δ1960 (**Figure 4B)**, consistent with structural studies showing that DPF2 interacts with a segment of the IDR2 domain (5).

**Figure 4.**
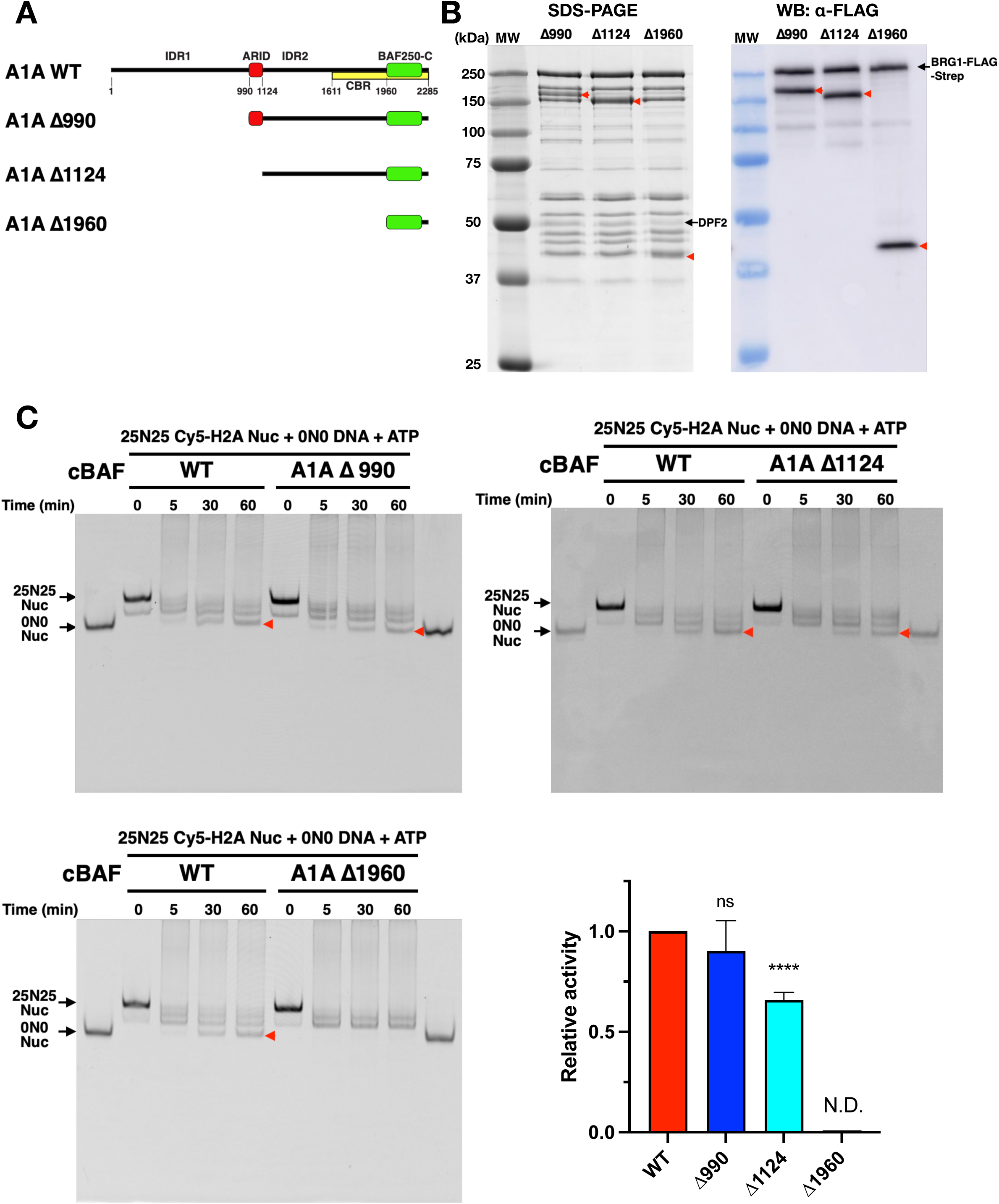
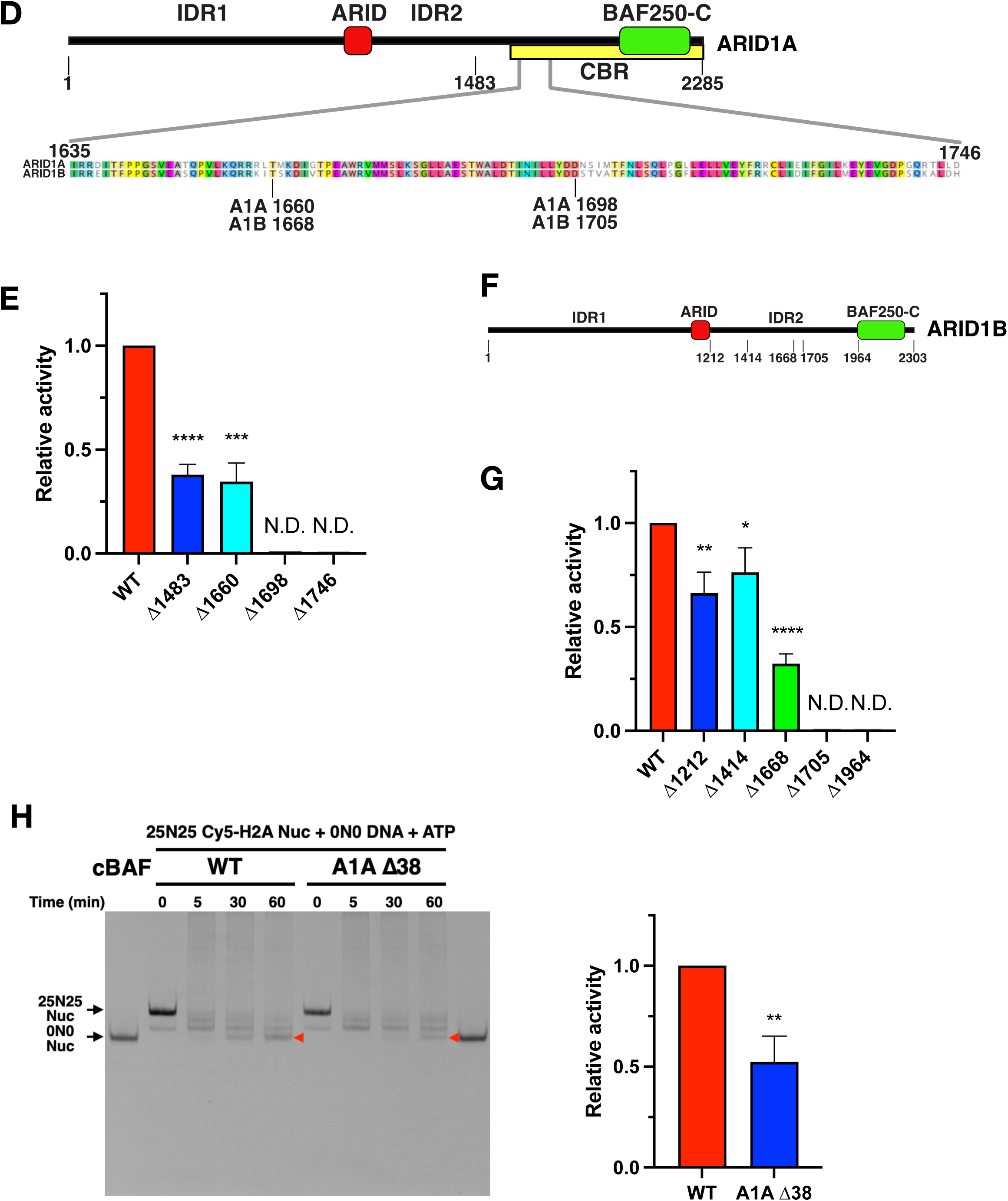
A part of IDR2 is required for histone transfer activity of cBAF. (A) Domain structure of ARID1A truncation constructs. CBR, core-binding region (ref (14)). (B) Coomassie-staining SDS-PAGE gel (Left panel) and western blot image (Right panel) of cBAF complexes containing ARID1A truncations. All ARID1A truncations and BRG1 contain 3xFLAG-tag at the N-terminus. (C) Representative Cy5-scanned native gel of histone octamer transfer assay for cBAF complexes containing ARID1A truncations. Red arrow indicates 0N0 nucleosomes produced by octamer transfer. Each activity was normalized to WT. (D) Sequence alignment of a region that is required for histone octamer transfer activity. (E) Quantification of histone octamer transfer activity of cBAF complexes containing ARID1A truncations. Each activity was normalized to WT. (F) Domain structure of ARID1B truncation constructs. (G) Quantification of histone octamer transfer activity of cBAF complexes containing ARID1B truncations. Each activity was normalized to WT. (H) Representative Cy5-scanned native gel and quantification of histone octamer transfer activity of cBAF complex containing ARID1A Δ38. The activity was normalized to WT. For Fig. 4C, 4E, 4G, and 4H, each error bar represents the standard error from at least three independent experiments using at least two independent BAF preparations. ****P<0.0001, ***P<0.001, **P<0.01, *P<0.05; ns, not significant; N.D., not detected.

Histone octamer transfer activity was measured for the cBAF complexes that harbor different ARID1A truncations. Removal of the IDR1 domain (A1A Δ990) had no significant impact on histone octamer transfer activity, whereas the additional removal of the ARID domain (A1A Δ1124) showed 30-40% reduction in activity (**Figure 4C**). Finally, removal of IDR1, IDR2, and the ARID domain (A1A Δ1960) eliminated histone octamer transfer activity (**Figure 4C**). These results indicated that ARID1A residues between 1124 and 1960 are required for histone transfer activity, though the ARID domain also appears to contribute.

Four additional ARID1A truncation derivatives were reconstituted into cBAF complexes, defining a ∼40 amino acid region that was required for optimal histone octamer transfer activity (compare A1A Δ1660 and A1A Δ1698; **Figures 4D and 4E, S4A and S4B**). This region is highly conserved between ARID1A and ARID1B, and it is one of a few short, structured domains within IDR2 (**Figure 4D**). Note that the A1A 1698 deletion eliminates histone octamer transfer activity without significant loss of the DPF2 subunit from cBAF (**Figure S4A**). We also generated various truncation constructs of ARID1B, and then reconstituted cBAF with each of these ARID1B variants. As for ARID1A, a similar, conserved region within ARID1B (a.a.1667-1705) was required for the histone octamer transfer activity of cBAF (**Figures 4F and 4G, S4C and S4D**). Finally, an internal deletion of ARID1A was constructed that removed only the 38 amino acid conserved domain. The cBAF complex harboring ARID1A ι138 retained only 52% of the wildtype level of histone octamer transfer activity, confirming the key role of this region (**Figures 4H and S4E**). These data also reinforce the view that other N-terminal domains of ARID1A, notably the ARID domain, also play a role in histone octamer transfer activity.

One possible mechanism of how this IDR2 region mediates the histone octamer transfer activity of cBAF is that the IDR2 region binds to a nucleosome and mediates histone octamer transfer. To test this possibility, we expressed and purified the entire IDR2 region **(Figure S4F**) to access its nucleosome binding activity. Electrophoretic mobility shift assays (EMSA) did not detect nucleosome binding activity for IDR2 **(Figure S4G**). In contrast, the protein fragment containing both the IDR2 and the ARID domain bound to nucleosomes, likely through the DNA binding of the ARID domain **(Figure S4H**).

Another possibility is that the IDR2 region has a nucleosome deposition activity like histone chaperones. To test this, the purified IDR2 region was incubated with Cy5-lebelled histone octamers and free 0N0 DNA fragments. The formation of 0N0 nucleosomes was not observed, suggesting that the IDR2 region does not have nucleosome deposition activity **(Figure S4I**). This data also confirmed that free histone octamers do not passively form nucleosomes onto DNA fragments under our assay conditions.

## DISCUSSION

Despite the clear importance of BAF complexes in development, transcriptional regulation, and disease prevention, distinct biological and biochemical functions for the 3 major forms of BAF complexes (cBAF, PBAF, and ncBAF) have remained largely unexplored. Recently a mononucleosome library screening approach found that cBAF, PBAF, and ncBAF complexes recognized distinct histone modification patterns (11). Here we reconstituted the cBAF, PBAF, and ncBAF complexes and directly compared their biochemical activities. Each complex showed distinct activities, with cBAF and ncBAF showing both higher affinity nucleosome binding and nucleosome sliding activity compared to PBAF. Perhaps not surprisingly, all three multi-subunit complexes were more active than the isolated BRG1 ATPase subunit. The trend of these biochemical properties is consistent with previous studies with endogenous BAF complexes (11).

In addition to reconstituting intact BAF complexes, we also probed how different subunits impact BAF assembly and activity. Largely consistent with previous work, we confirmed that the BAF170/BAF155 subunits are key for assembly of cBAF. Furthermore, our biochemical analysis of BAF8 subcomplexes suggested that the Base module is important for both nucleosome-binding and sliding activities, whereas the ARP module is required only for robust levels of nucleosome sliding, consistent with the notion that the ARP module is involved in “coupling” of the ATPase and sliding activities (15). The most striking result from this work was our finding that ARID1A was required for the histone octamer transfer activity of cBAF. Loss of ARID1A from cBAF had little impact on nucleosome binding affinity or nucleosome sliding activity, indicating that it provides a unique function. Since both PBAF and ncBAF complexes also exhibit histone octamer transfer activity, our data suggests that complex-specific subunits, such as ARID2 and GLTSCR1, may play a similar role.

The mechanism by which ARID1A/ARID1B mediates the histone octamer transfer activity of cBAF is unclear. Our analysis of a series of ARID1A truncations suggests that a 38-amino acid domain of ARID1A is required for the histone octamer transfer activity of cBAF. Interestingly, in the cryoEM structure of cBAF, this domain is comprised of 2 alpha-helixes at the end of the ARM-like structure of the C-terminal domain of ARID1A. C-terminal to this domain, the ARID1A polypeptide extends to the proximity of the DPF2 subunit, returning to form the remainder of ARM-like structure. One possibility is that this new functional domain mediates the transfer of histone octamers directly through an association with either histones or DNA. Consistent with this model, our deletion analyses indicates that this domain may function in concert with the ARID DNA binding domain. Given that this region is near the ARM domain that interacts with the BRG1 scaffold, an alternative view is that deletion of this domain may alter the conformation of the Base module which may be key for mediating histone octamer transfer activity.

Recent genome-wide localization studies indicate that BAF complexes occupy nearly 41,000 genomic sites that include TSS-proximal (e.g. promoters) and TSS-distal (e.g. enhancers) locations (14). These localization patterns are consistent with the established role for BAF complexes in broadly controlling enhancer function (16–19). Loss of ARID1 subunits disrupts the recruitment of BAF complexes to ∼10,000 of these genomic sites, primarily impacting TSS-distal locations (14). Furthermore, loss of ARID1 subunits reduces chromatin accessibility at many sites (14,17,20,21). ARID1A domain mapping studies indicated that N-terminal regions of ARID1A play key roles in BAF targeting and recruitment (14). Although much emphasis has been placed on the IDR1 region which drives phase condensation in vitro and in vivo, regions that we identify here as important for histone octamer transfer activity are also encompassed by deletions that impair BAF targeting and in vivo activity (14). Given that BAF targeting is also associated with increased chromatin accessibility, it seems likely that histone octamer transfer activity may play a dominant role in facilitating the formation of open chromatin domains.

## DATA AVAILABILITY

The data are available from the corresponding author upon reasonable request.

## ACKNOWLEDGEMENTS

We thank Craig L. Peterson and members of the Peterson Lab for their constructive discourse.

## AUTHOR CONTRIBUTIONS

Naoe Moro (Investigation, Formal analysis, Resources, Methodology, Data curation, Writing-review & editing)

Yukiko Fujisawa-Tanaka (Resources, Methodology, Writing-review & editing)

Shinya Watanabe (Conceptualization, Data curation, Funding acquisition, Investigation, Methodology, Supervision, Validation, Visualization, Writing-original draft, Writing-review & editing)

## MATERIAL and METHODS

### Insect cell lines

Sf9 cells (Thermo Fisher Scientific, Waltham, MA, USA) were used for baculovirus production and recombinant protein expression, and grown in ESF921 media (Expression Systems, Davis, CA, USA) at 27 °C.

### Expression and purification of human BAF complexes

Human BAF subunit genes were cloned and expressed using the MultiBac baculovirus expression system (Geneva Biotech, Geneva, Switzerland). Genes coding for BRG1 (residue 1– 1647 with an N-terminal 3× FLAG-tag and a C-terminal twin Strep-tag), GLTSCR1(residue 1– 1560 with a C-terminal twin Strep-tag), ARID1A, ARID1B, ARID2, β-Actin, BAF53a, BCL7a, DPF2, BRD7, BRD9, PBRM1, PHF10, SMARCB1, SMARCC1, SMARCC2, SMARCD1, SMARCE1, and SS18 were synthesized with codon optimization for insect cells (Genewiz, Cambridge, MA, USA). For cBAF, 12 subunits were combined in 2 separate bacmids. For PBAF, 13 subunits were combined in one bacmid. For ncBAF, 10 subunits were combined in 2 separate bacmids. For BAF8 and BAF10, 8 and 10 subunits were combined in one bacmid, respectively. Sf9 cells were infected with viruses and cultured for 72 h at 27 °C. Cells were harvested and washed with phosphate-buffered saline and then lysed by sonication in Lysis buffer (350 mM NaCl, 20 mM HEPES [pH 7.5], 10% glycerol, 0.1% Tween, 1 mM MgCl_2_, 50 µM ZnCl_2_, 1 mM DTT, 1 mM phenylmethylsulphonyl fluoride, 1 mM benzamidine, 2 µg/mL Leupeptin, 2 µg/mL Pepstatin A, and 2 µg/mL Chymostatin [Sigma-Aldrich, St. Louis, MO, USA]). After centrifugation at 40 000 *g* and 4 °C for 20 min, the supernatant was loaded onto a StrepTactin HP column (Cytiva, Marlborough, MA, USA). The column was washed with Lysis buffer and Buffer A (150 mM NaCl, 20 mM HEPES [pH 7.5], 10% glycerol, 1 mM MgCl_2_, and 1 mM DTT) and eluted with Buffer A plus 10 mM desthiobiotin (Sigma-Aldrich). The eluted proteins were then subjected to a Q Sepharose column (Cytiva). After washing with Buffer A, the proteins were eluted by a linear salt gradient. The peak fractions were concentrated in buffer A and flash-frozen in liquid nitrogen. Protein concentration was determined by the densitometry analysis of SDS-PAGE using BSA as a standard. The molar concentration of each BAF subcomplex was estimated from the BRG1 subunit concentration. For PBAF and ncBAF, the molar concentration was normalized based on their ATPase activities compared to cBAF with known concentration. Subunit compositions were confirmed by SDS-PAGE and mass spectrometry.

### Nucleosome preparation

Recombinant human histones were expressed in *Escherichia coli* cells and purified as previously described (22,23). In brief, expressed histones were purified as inclusion bodies, solubilized in unfolding buffer (7 M guanidinium hydrochloride, 20 mM Tris-HCl [pH 7.5], and 10 mM DTT), and dialyzed against urea dialysis buffer (7 M urea, 10 mM Tris-HCl [pH 8.0], 0.1 M NaCl, 1 mM EDTA, 0.2 mM phenylmethylsulphonyl fluoride, and 5 mM 2-mercaptoethanol). Samples were injected into tandemly connected Q Sepharose and SP Sepharose columns, and eluted from SP Sepharose by a linear salt gradient. Histone fractions were dialyzed against water with 0.2 mM phenylmethylsulphonyl fluoride and 5 mM 2-mercaptoethanol, and lyophilized. Histone H2A (T120C) were labelled with Cy5, as previously described (24). The four histones (H2A, H2B, H3, and H4) were mixed in equimolar ratios in unfolding buffer, dialyzed against refolding buffer (2 M NaCl, 10 mM Tris-HCl [pH 7.5], 1 mM EDTA, and 5 mM 2-mercaptoethanol) and purified through a Superdex-200 column. Nucleosomes were reconstituted by mixing octamers with DNA at a 1:1 ratio in HI buffer (2 M NaCl, 10 mM Tris-HCl [pH 7.5], and 5 mM 2-mercaptoethanol), and dialyzed against a linear salt gradient buffer from HI to LO buffer (50 mM NaCl, 10 mM Tris-HCl [pH 7.5], and 5 mM 2-mercaptoethanol) for 20 h.

### Histone octamer transfer assays

For histone octamer transfer assays, Cy5-labelled histone H2A-containing mononucleosomes were reconstituted by salt dialysis onto 197 bp DNA fragments containing the 601 nucleosome-positioning sequence in the center of the DNA fragment (25N25). Mononucleosomes (15 nM) were incubated with BAF (60 nM), 0N0 DNA fragments (10 nM), and 2 mM ATP in Buffer B (25 mM NaCl, 25 mM HEPES [pH 7.5], 5 mM MgCl_2_, and 1 mM DTT) at 30 °C for 60 min. The reactions were initiated by addition of ATP. At each time point, the reactions were quenched with 5% glycerol and 0.1 mg/ml salmon sperm DNA, incubated for 5 min at 30 °C, and resolved on 5% Native-PAGE in 0.5 X TBE. Gels were scanned with Cy5 using a Typhoon biomolecular imager (GE Healthcare, Chicago, IL, USA). Biotinylated 25N25 DNA fragments were generated by PCR using biotinylated primers. The fraction of 0N0 nucleosome product was quantified and plotted over time. Transfer rate obtained from the plot was normalized to a rate of the wild-type cBAF.

For FRET assays, Cy5-labelled histone H2A-containing mononucleosomes were reconstituted by salt dialysis onto 25N25 DNA fragments. Mononucleosomes (15 nM) were incubated with BAF (60 nM), Cy3-labelled 0N0 DNA fragments (10 nM) and 2 mM ATP in buffer B at 30 °C for 60 min. The FRET signal was monitored using a Tecan Spark microplate reader (Tecan, Switzerland) with excitation at 530 nm and emission at 670 nm.

### Nucleosome sliding assays

*HhaI* restriction enzyme accessibility assays were performed as described (25). Mononucleosomes were reconstituted by salt dialysis onto Cy5-lablelled 25N25 DNA fragments. Mononucleosomes (15 nM) were incubated with BAF (2nM, 3nM, or 16nM), 4 units/µl *HhaI*, and 2 mM ATP in buffer B at 30 °C. At each time point, the reactions were stopped by addition of STOP buffer (10 mM HEPES [pH 7.5], 40 mM EDTA, 0.6% SDS, 5% glycerol, 0.1 mg/ml Proteinase K), incubated for 20 min at 50 °C, and resolved on 6% Native-PAGE in 0.5 X TBE. Gels were scanned with Cy5 using a Typhoon biomolecular imager. The fraction of cut DNA was quantified and plotted over time. Sliding rate was obtained from an initial rate of the plot.

### Fluorescence polarization (FP) assays

Fluorescence polarization assays were performed as described (26). Various concentrations of BAF were incubated with 10 nM Cy5-labelled mononucleosomes in Buffer B, except for 60 mM NaCl. FP were measured using a Tecan Spark microplate reader. All binding curves were fit to the quadratic binding equation:

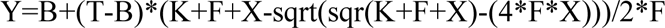

where B and T are the minimum and maximum FP signals, respectively, K is the apparent dissociation constant, F is the nucleosome concentration, and X is the concentration of BAF.

### ATPase assays

ATPase assays were performed as previously described (27). ATP hydrolysis was monitored using NAD/NADH-coupled ATP regeneration system with lactate dehydrogenase (LDH) and pyruvate kinase (PK). 50 nM BAF was incubated with 10% PK/LDH mixture (Sigma), 1 mM phosphoenolpyruvate, 1 mM NADH, and 1 mM ATP in Buffer B. 100 nM 25N25 DNA fragment was used as a substrate. NADH absorbance was measured at 340 nm using a Tecan Spark microplate reader. ATPase activity was obtained from an initial linear rate subtracted from background signal (without BAF).

### Histone eviction assays

For histone eviction assays, mononucleosomes were reconstituted by salt dialysis onto Cy5-labelled 25N25 DNA fragments. Mononucleosomes (15 nM) were incubated with BAF (60 nM) and 2 mM ATP in Buffer B at 30 °C for 90 min. The reactions were initiated by addition of ATP. At each time point, the reactions were quenched with 5% glycerol and 0.1 mg/ml salmon sperm DNA, incubated for 5 min at 30 °C, and resolved on 5% Native-PAGE in 0.5 X TBE. Gels were scanned with Cy5 using a Typhoon imager. The fraction of free 25N25 DNA was quantified and plotted over time.

### Electrophoretic mobility shift assays (EMSA) for nucleosome binding and deposition

For nucleosome binding assays, Cy5-labelled histone H2A-containing mononucleosomes were reconstituted by salt dialysis onto 0N0 DNA fragments. Mononucleosomes (15 nM) were incubated with various concentrations of IDR2 or ARID+IDR2 in Buffer B for 15 min at room temperature. The reactions were resolved on 5% Native-PAGE in 0.25 X TBE. Gels were scanned using a Typhoon imager.

For nucleosome deposition assays, Cy5-labelled histone H2A-containing octamers and 0N0 DNA fragments were mixed in equimolar ratios (each 100 nM) with various concentrations of IDR2 in Buffer B for 15 min at room temperature. The reactions were resolved on 5% Native-PAGE in 0.5 X TBE. Gels were scanned using a Typhoon imager.

