## Supplemental Figures for "ARID1A regulates histone octamer transfer activity of human canonical BAF complex"

### Supplemental Figure S1

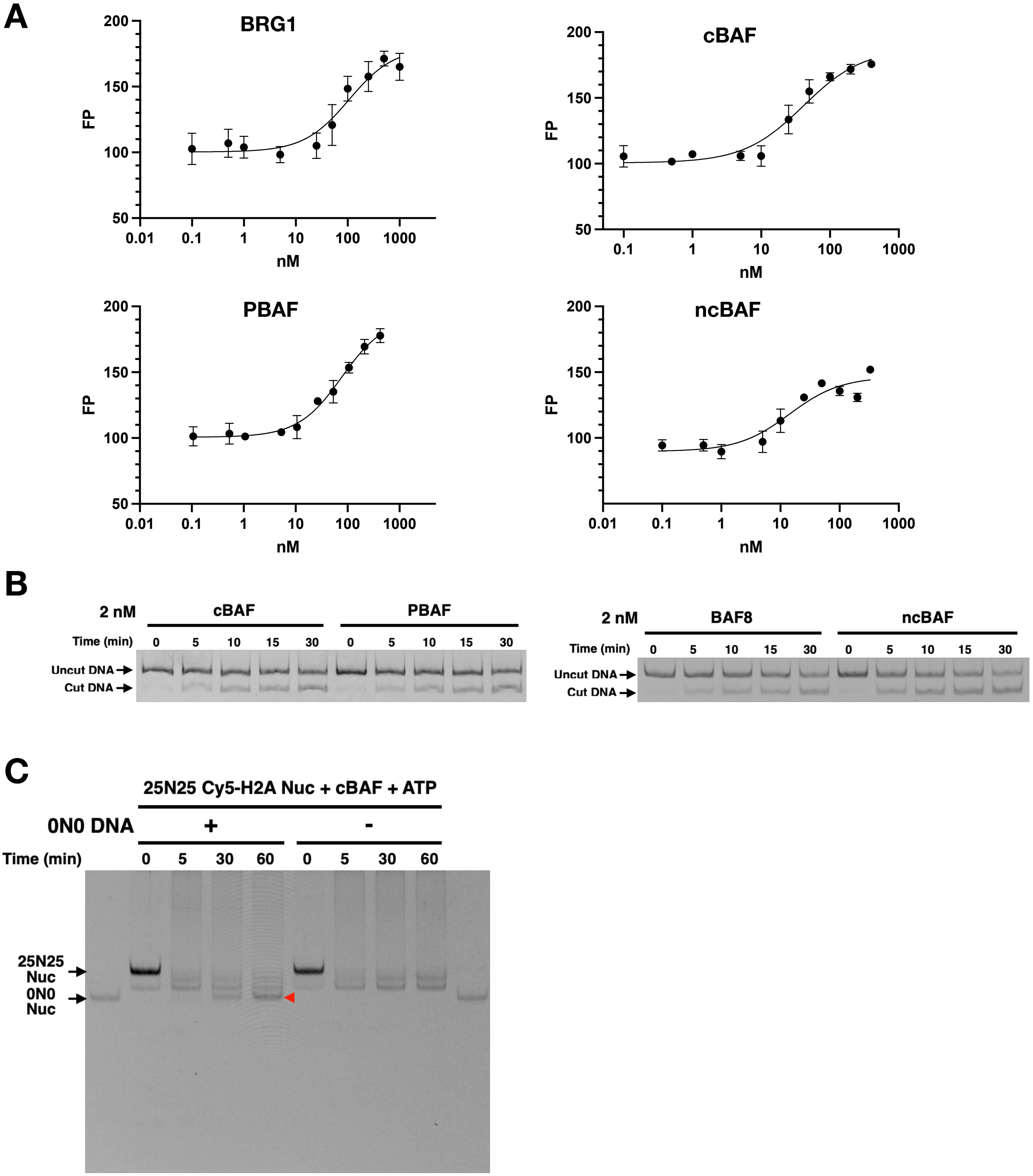

**D**

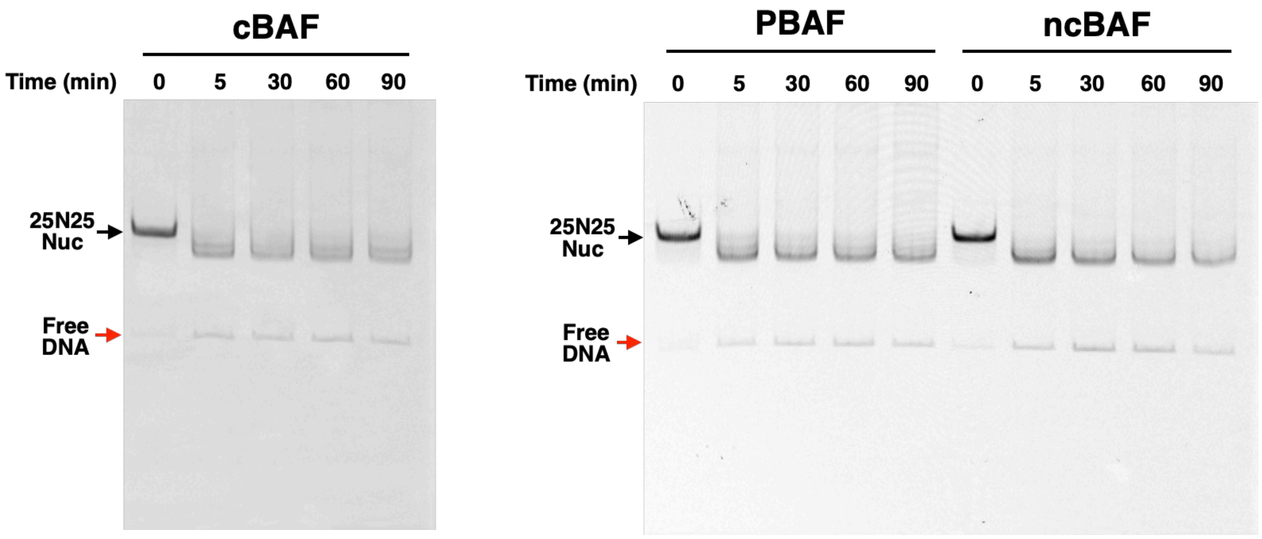
cBAF
Time (min) 0 5 30 60 90
25N25 Nuc
Free DNA
PBAF
ncBAF
Time (min) 0 5 30 60 90 0 5 30 60 90
25N25 Nuc
Free DNA

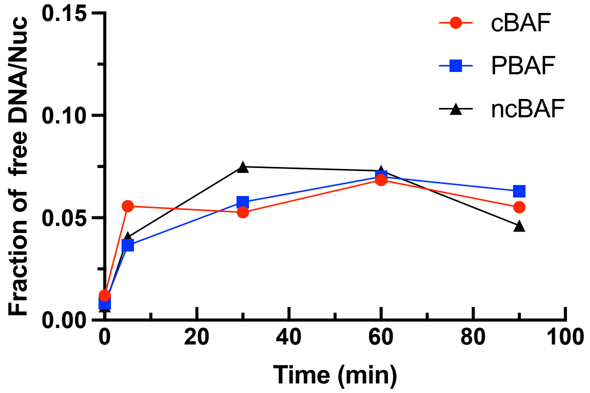
Fraction of free DNA/Nuc
0.15
0.10
0.05
0.00
0
20
40
60
80
100
Time (min)
cBAF
PBAF
ncBAF

**Figure S1. Functional analyses of cBAF, PBAF, and ncBAF, related to Figure 1.**

(A) Representative FP assay for BRG1, cBAF, PBAF, and ncBAF with 10 nM Cy5-labelled nucleosomes. Kd and SE were calculated from at least three replicates.

(B) Representative Cy5-scanned native gels of HhaI-restriction enzyme accessibility assay for cBAF, PBAF, and ncBAF. Sliding rate and SE were determined from an initial rate of sliding with at least three replicates.

(C) Representative Cy5-scanned native gels of histone octamer transfer assay in the presence or absence of the 0N0 DNA fragment as an acceptor. Red arrow indicates 0N0 nucleosomes formed by octamer transfer.

(D) Representative Cy5-scanned native gels and quantification of histone eviction assay. Fractions of free 0N0 DNA were quantified and plotted over time. Each plot was an average from duplicate.

### Supplemental Figure S2

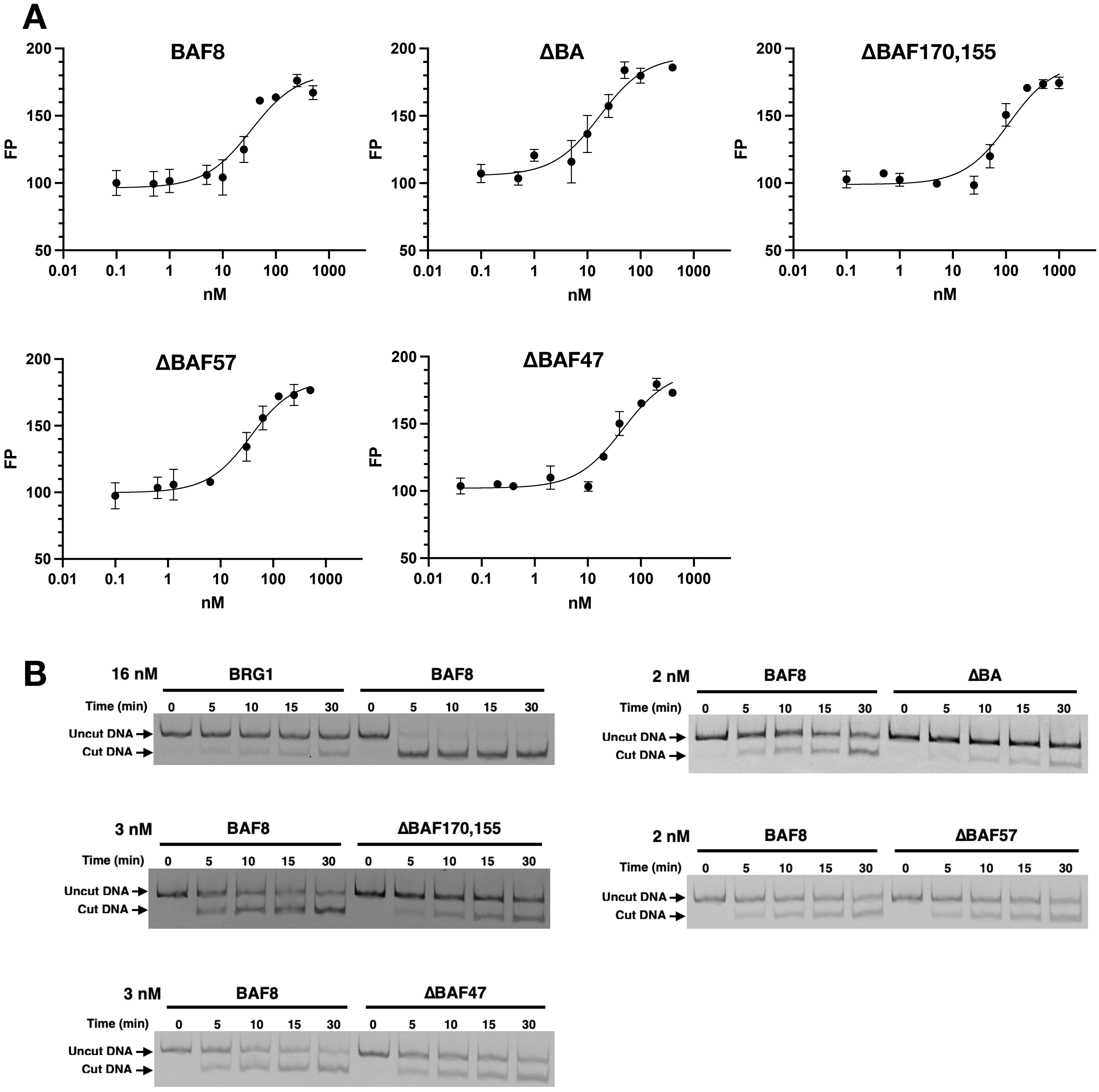

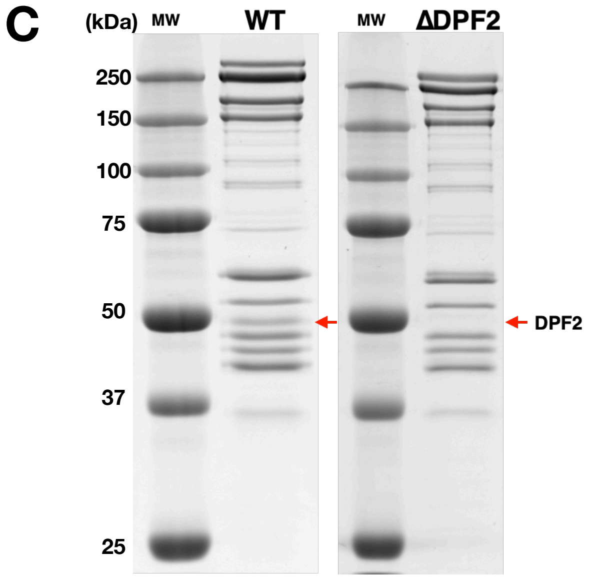

C
(kDa)
MW
WT
MW
ΔDPF2
250
150
100
75
50
37
25
DPF2

**Figure S2. Functional analyses of BAF8 subcomplexes, related to Figure 2.**

(A) Representative FP assay for BAF8, ΔBA, ΔBAF170,155, ΔBAF57, and ΔBAF47 with 10 nM Cy5-labelled nucleosomes. Kd and SE were calculated from at least three replicates.

(B) Representative Cy5-scanned native gels of HhaI-restriction enzyme accessibility assay for BRG1, BAF8, ΔBA, ΔBAF170,155, ΔBAF57, and ΔBAF47. Sliding rate and SE were determined from an initial rate of sliding with at least three replicates.

(C) Coomassie-staining SDS-PAGE gels of purified cBAF WT and ΔDPF2 complexes.

### Supplemental Figure S3

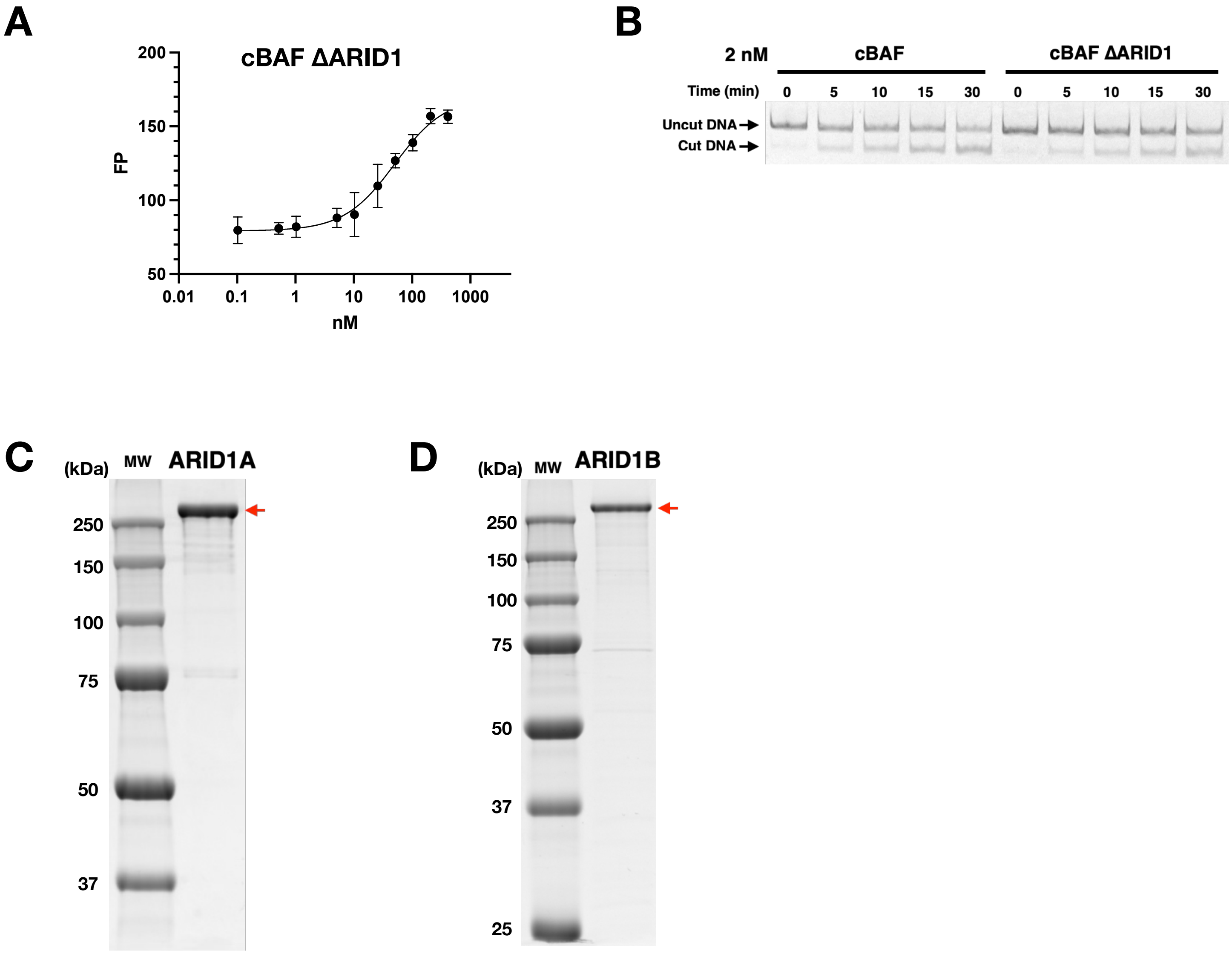

**Figure S3. ARID1A/ARID1B are required for histone octamer transfer activity, related to Figure 3.**

(A) Representative FP assay for cBAF ΔARID1 with 10 nM Cy5-labelled nucleosomes. Kd and SE were calculated from at least three replicates.

(B) Representative Cy5-scanned native gel of HhaI-restriction enzyme accessibility assay for cBAF ΔARID1. Sliding rate and SE were determined from an initial rate of sliding with at least three replicates.

(C) Coomassie-staining SDS-PAGE gel of purified recombinant ARID1A.

(D) Coomassie-staining SDS-PAGE gel of purified recombinant ARID1B.

### Supplemental Figure S4

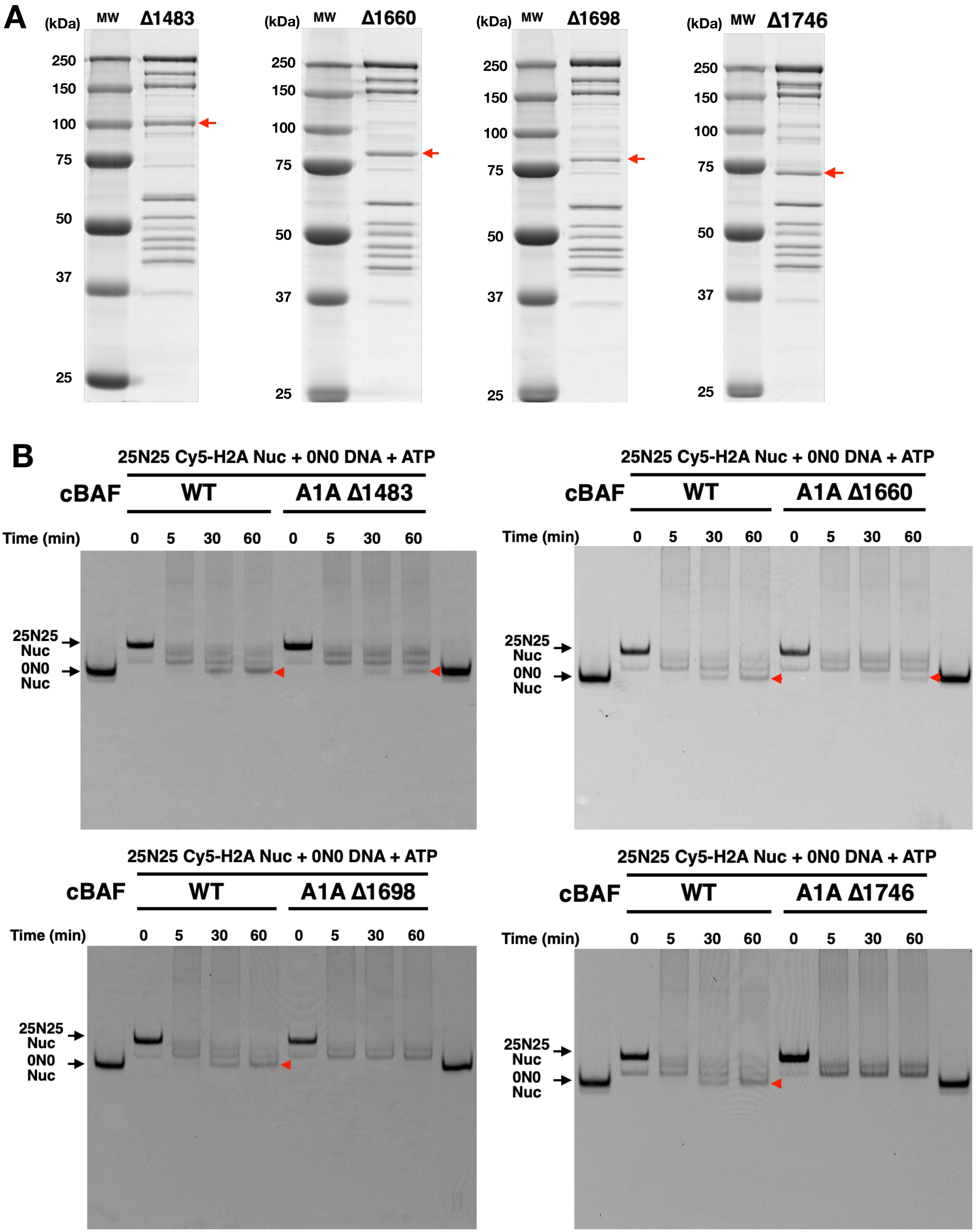

C

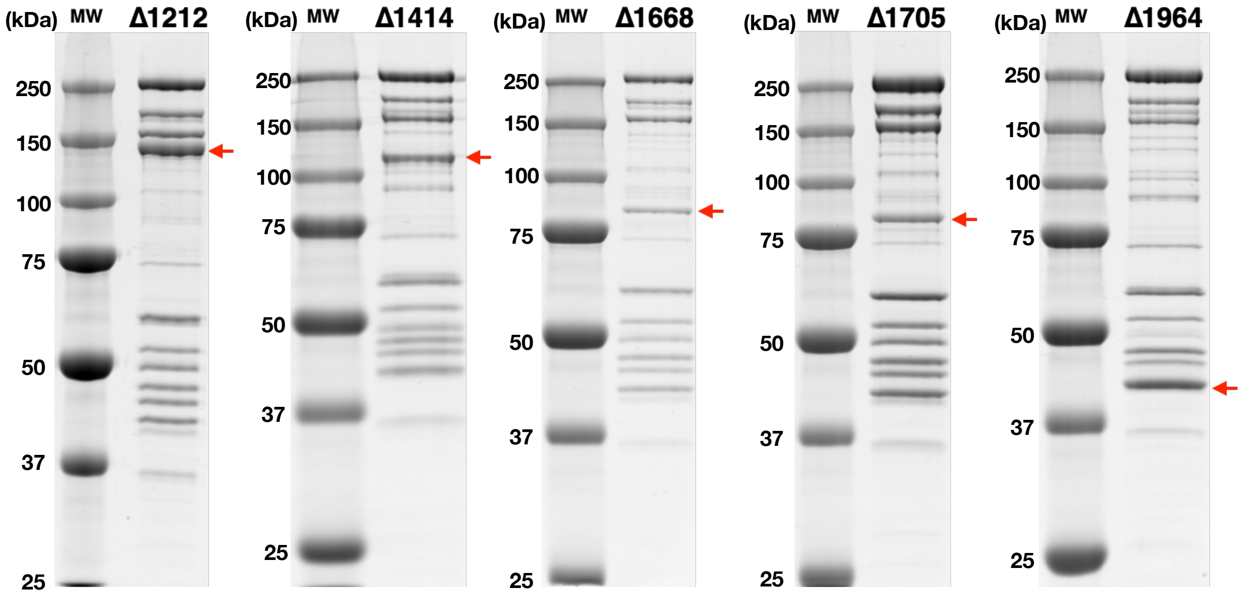

(kDa) MW Δ1212
250
150
100
75
50
37
25
(kDa) MW Δ1414
250
150
100
75
50
37
25
(kDa) MW Δ1668
250
150
100
75
50
37
25
(kDa) MW Δ1705
250
150
100
75
50
37
25
(kDa) MW Δ1964
250
150
100
75
50
37
25


D

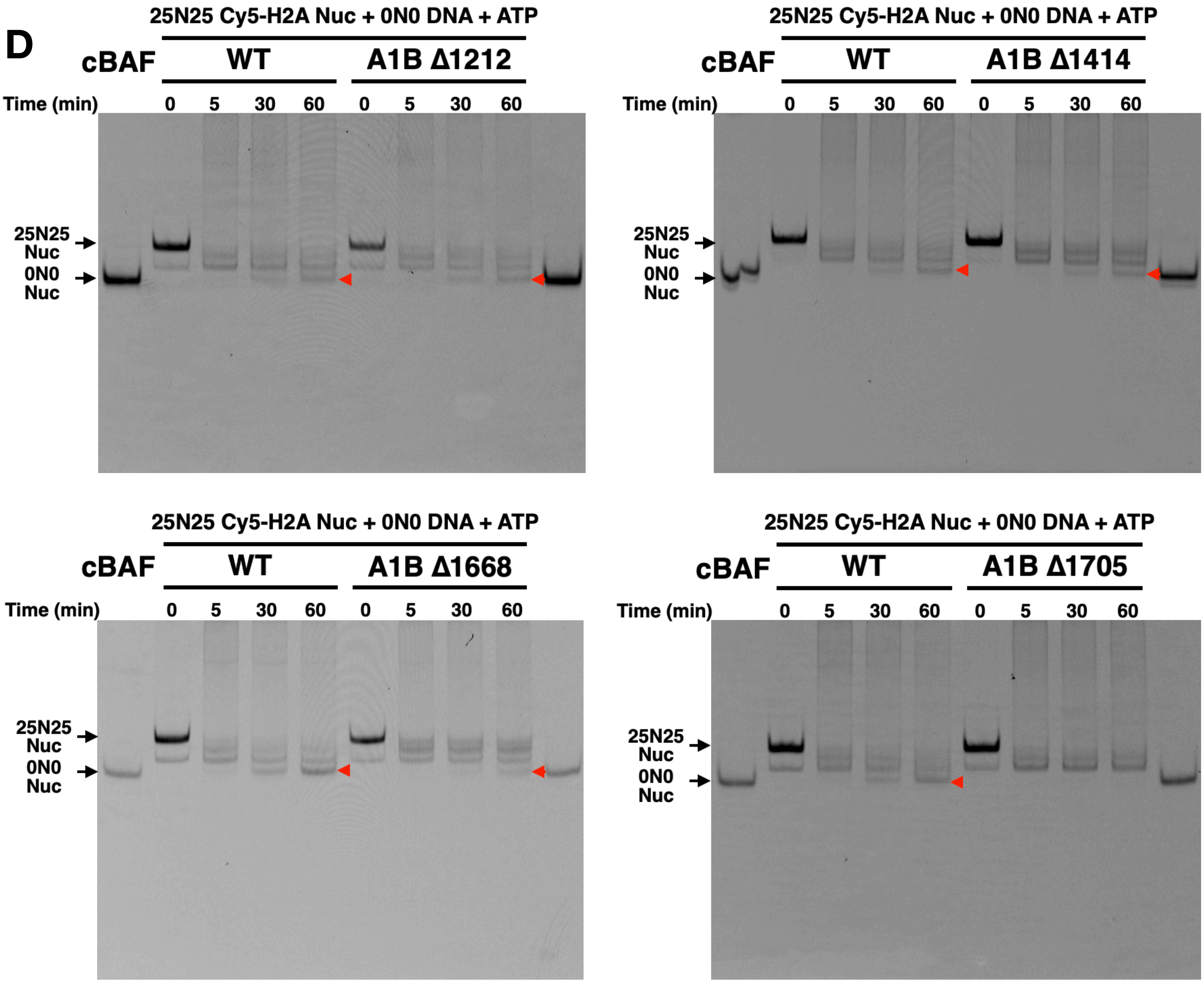

25N25 Cy5-H2A Nuc + 0N0 DNA + ATP
cBAF WT A1B Δ1212
Time (min) 0 5 30 60 0 5 30 60
25N25 Nuc
0N0 Nuc
25N25 Cy5-H2A Nuc + 0N0 DNA + ATP
cBAF WT A1B Δ1414
Time (min) 0 5 30 60 0 5 30 60
25N25 Nuc
0N0 Nuc
25N25 Cy5-H2A Nuc + 0N0 DNA + ATP
cBAF WT A1B Δ1668
Time (min) 0 5 30 60 0 5 30 60
25N25 Nuc
0N0 Nuc
25N25 Cy5-H2A Nuc + 0N0 DNA + ATP
cBAF WT A1B Δ1705
Time (min) 0 5 30 60 0 5 30 60
25N25 Nuc
0N0 Nuc

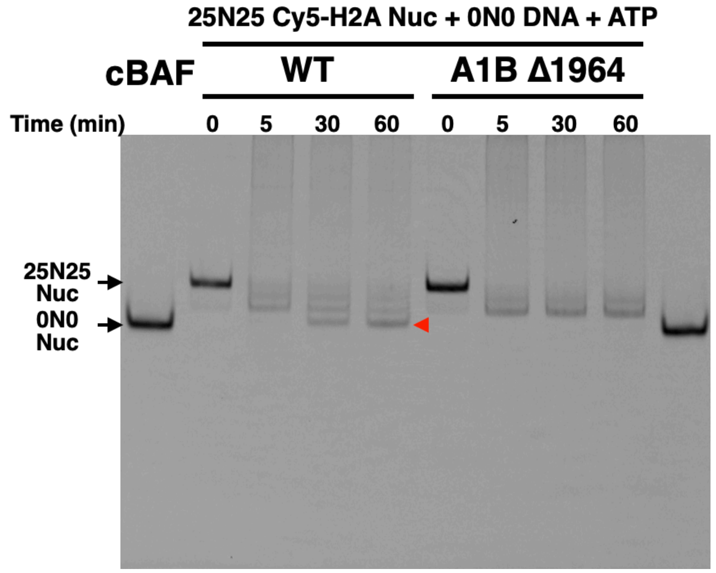

25N25 Cy5-H2A Nuc + 0N0 DNA + ATP
cBAF
WT
A1B Δ1964
Time (min)
0
5
30
60
0
5
30
60
25N25 Nuc
0N0 Nuc


E

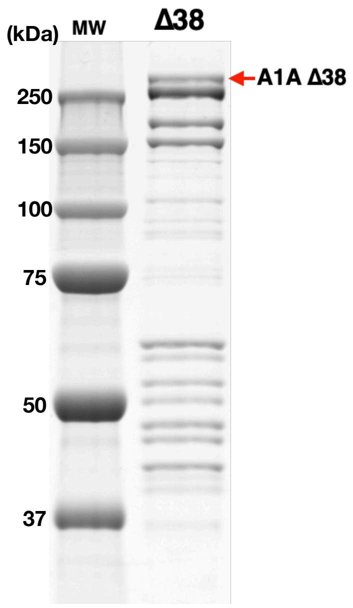

(kDa)
MW
Δ38
250
150
100
75
50
37
A1A Δ38


F

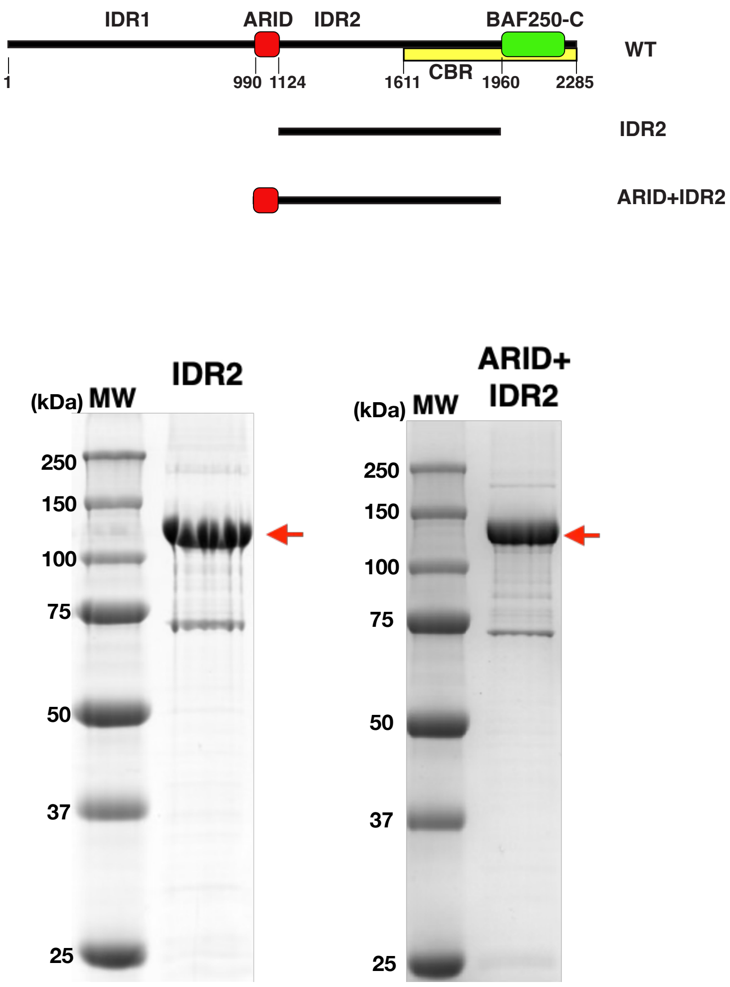

IDR1
ARID
IDR2
BAF250-C
WT
1
990
1124
1611
CBR
1960
2285
IDR2
ARID+IDR2
(kDa) MW
IDR2
250
150
100
75
50
37
25
(kDa) MW
ARID+ IDR2
250
150
100
75
50
37
25

G

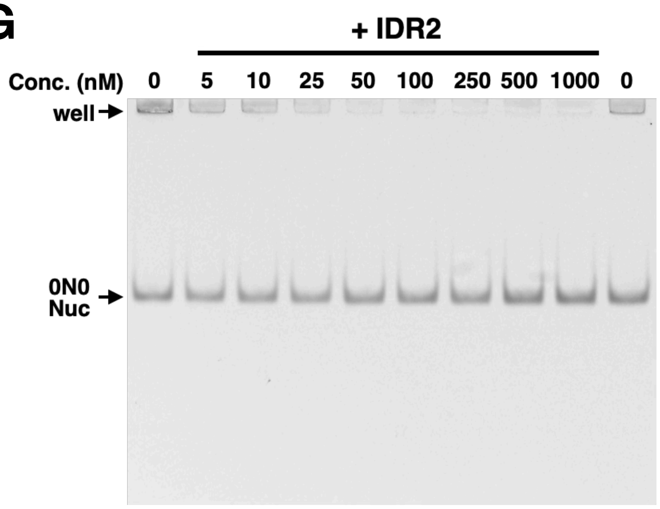
+ IDR2
Conc. (nM)
0
5
10
25
50
100
250
500
1000
0
well
0N0
Nuc

I

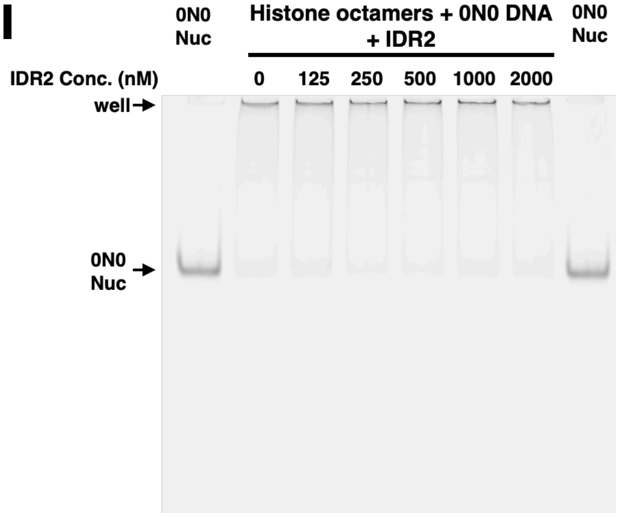
0N0
Nuc
Histone octamers + 0N0 DNA
+ IDR2
0N0
Nuc
IDR2 Conc. (nM)
0
125
250
500
1000
2000
well
0N0
Nuc

H

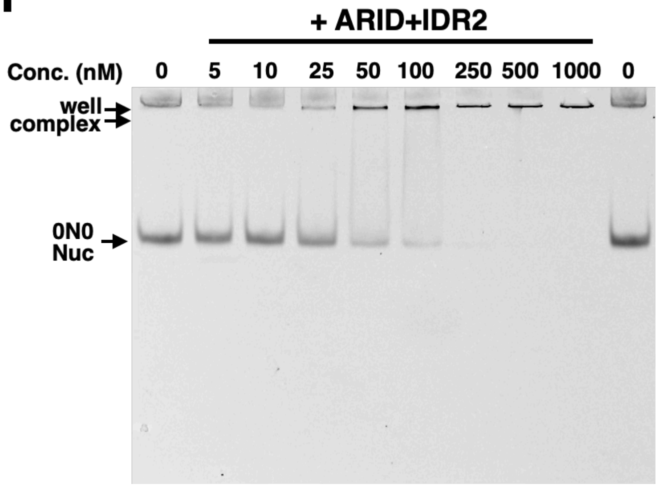
+ ARID+IDR2
Conc. (nM)
0
5
10
25
50
100
250
500
1000
0
well
complex
0N0
Nuc

**Figure S4. Histone octamer transfer activity of cBAF containing ARID1A or ARID1B truncations, related to Figure 4.**

(A) Coomassie-staining SDS-PAGE gels of cBAF containing ARID1A Δ1483, Δ1660, Δ1698, and Δ1746. Red arrows indicate ARID1A truncations.

(B) Representative Cy5-scanned native gels of histone octamer transfer assay for cBAF containing ARID1A Δ1483, Δ1660, Δ1698, and Δ1746. Red arrows indicate 0N0 nucleosomes produced by octamer transfer. Relative activity and SE were calculated from at least three replicates.

(C) Coomassie-staining SDS-PAGE gels of cBAF containing ARID1B Δ1212, Δ1414, Δ1668, Δ1705 and Δ1964. Red arrows indicate ARID1B truncations.

(D) Representative Cy5-scanned native gels of histone octamer transfer assay for cBAF containing ARID1B Δ1212, Δ1414, Δ1668, Δ1705 and Δ1964. Red arrows indicate 0N0 nucleosomes produced by octamer transfer. Relative activity and SE were calculated from at least three replicates.

(E) Coomassie-staining SDS-PAGE gel of purified cBAF containing ARID1A Δ38.

(F) Domain structure of IDR2 constructs. Coomassie-staining SDS-PAGE gels of purified recombinant IDR2 and ARID+IDR2.

(G, H) Representative Cy5-scanned native gels of nucleosome-binding of IDR2 (G) or ARID+IDR2 (H). Mononucleosomes (15 nM) were incubated with the indicated amount of IDR2 or ARID2+IDR2.

(I) Representative Cy5-scanned native gel of nucleosome deposition assays for IDR2. Cy5-labelled octamers and 0N0 DNA were mixed in equimolar ratios (each 100 nM) with the indicated amount of IDR2. 0N0 nucleosomes (100 nM) were loaded as a control.

**Table S1. Mass spectrometry data for purified cBAF complex.**

Spectrum counts on cBAF subunits. Note that each of cBAF, PBAF, and ncBAF-specific gel bands was also confirmed by mass spectrometry.

### Supplemental Table S1

| cBAF subunit | Spectrum Count |
|---|---|
| BRG1 | 591 |
| ARID1A | 577 |
| BAF170 (SMARCC2) | 307 |
| BAF155 (SMARCC1) | 346 |
| BAF60 (SMARCD1) | 328 |
| BAF57 (SMARCE1) | 67 |
| BAF47 (SMARCB1) | 65 |
| BAF53a (ACTL6A) | 91 |
| ß-Actin | 97 |
| DPF2 | 69 |
| SS18 | 20 |
| BCL7a | 82 |
